# Phylogenomics and Biogeography of North American Trechine Cave Beetles (Coleoptera: Carabidae)

**DOI:** 10.1101/2023.04.27.538603

**Authors:** Joseph B. Benito, Karen A. Ober, Matthew L. Niemiller, Karen A. Ober

## Abstract

Cave trechines beetles (Coleoptera: Carabidae: Trechini) are members of cave communities globally and important models for understanding the colonization of caves, adaptation to cave life, and the diversification of cave-adapted lineages. In eastern North America, cave trechines are the most species-rich group of terrestrial troglobionts, comprised of over 150 taxa in six genera with no extant surface members. Previous studies have hypothesized the climate change during the Pleistocene was a major driver of cave colonization and diversification in this and other temperate terrestrial cave fauna. However, our time-calibrated molecular phylogeny resulting from the analysis of 16,794 base pairs (bp) from 68 UCE loci for 45 species of the clade supports an alternative hypothesis whereby cave colonization of the surface ancestor of eastern North American cave trechines likely began in the middle Miocene in the Appalachians Ridge and Valley (APP) and dispersed into the Interior Low Plateau (ILP) in an east to west manner around 11.5 Mya. The APP served as a cradle for diversification and also as a bridge linking the southern Appalachians and Interior Low Plateau enabling the dispersal and subsequent diversification of these cave beetles. Major clades in our time-calibrated phylogeny attained their present-day geographic distributions by the early Miocene followed by multiple additional episodes of cave colonization and diversification occurring throughout the Pliocene and Pleistocene. The genera *Neaphanops*, *Darlingtonea*, *Nelsonites*, and *Ameroduvalius* were nested within specious genus *Pseudanopthalmus* supporting the hypothesis that these genera are derived Pseudanophtlamus. Moreover, while several morphologically-derived species groups of *Pseudanopthalmus* were recovered as monophyletic, others were not warranting future taxonomic and systematic research. The molecular systematics and biogeography of these unique cave beetles offer a model for other comparative evolutionary and ecological studies of troglobionts to further our understanding of factors driving speciation and biogeographic patterns.

## Introduction

Caves are home to unique and diverse communities, and they represent one of the most unforgiving environments on Earth. The primary characteristic of caves and subterranean habitats is the lack of light and associated photosynthesis leading to limited food resources (Culver and Pipan 2009; Soares and Niemiller 2020). Cave organisms tend to exhibit similar morphological, physiological, and behavioral traits, such as loss of eyes, reduction of pigmentation, development of nonvisual sensory organs, changes in metabolism, and longer lifespan (termed troglomorphy; Racovitza 1907; Vandel 1964; Culver et al. 1990; Culver and Pipan 2009). Caves and their fauna have been viewed as model systems for addressing fundamental questions in evolutionary biology, ecology, biogeography, and speciation (Poulson and White 1969; Juan et al. 2010; Mammola 2019). However, testing hypotheses related to speciation, evolution, and biogeography of cave fauna often has been hampered in the past due to difficulties associated with sampling subterranean habitats, convergence in morphology, and extinction of surface ancestors (Holsinger 2000; Porter 2007; Juan and Emerson 2010). Despite these challenges, new molecular approaches developed in recent years have provided an opportunity to better understand the evolutionary and biogeographic processes that facilitate adaptation and speciation and shape distributions in subterranean environments (Jeffery 2009; Juan et al. 2010; Liu et al. 2017; Torres-Dowdall et al. 2018). Yet, many questions remain unanswered or poorly investigated (Juan et al. 2010; Morvan et al. 2013; Mammola et al. 2019). For example, the relationships between many subterranean and surface lineages remain largely unknown and the timing and patterns of diversification of many cave organisms are understudied.

In North America, subterranean biodiversity is primarily associated with distinct karst regions (Culver et al. 2003; Hobbs 2012; Niemiller et al. 2019), a type of landscape underlain by carbonate rocks that possess dense networks of subterranean drainage systems, sinkholes, springs, and caves (Ford and Williams 2007). Four karst regions – Interior Low Plateau, Appalachians, Edwards Plateau & Balcones Escarpment, and Ozarks – account for nearly 80% of the more than 1350 described troglobiotic (terrestrial) and stygobiotic (aquatic) diversity in the continental United States and Canada (Niemiller et al. 2019). Several phylogenetic studies in recent years have advanced our knowledge of the biogeography, evolution, and systematics of many subterranean organisms in North America, shedding light on distributional patterns, cryptic diversity, colonization history, and modes of speciation. However, most studies have focused largely on three primarily aquatic taxonomic groups that account for only ∼4% of subterranean biodiversity in the United States and Canada (Niemiller et al. 2019): cavefishes (e.g., Dillman et al. 2011; Garcia-Machado et al. 2011; Niemiller et al. 2012, 2013a,b; Strecker et al. 2012; Fumey et al. 2018; Hart et al. 2020), salamanders (e.g., Chippindale et al. 2000; Hillis et al. 2001; Wiens et al. 2003; Niemiller et al. 2008, 2009; Bendik et al. 2013; Phillips et al. 2017; Devitt et al. 2019; Grant et al. 2022), and crayfishes (e.g., Buhay and Crandall 2005, 2008, 2009; Buhay et al. 2007; Stern et al. 2017; Dooley et al. 2022). Recent North American phylogenetic studies involving troglobiotic invertebrates are comparatively few, but include spiders (e.g., Hedin 1997a,b, 2015; Paquin and Hedin 2004; Snowman et al. 2010; Hedin et al. 2018), harvestmen (Derkarabetian et al. 2010, 2022; Hedin and Thomas 2010), springtails (Katz et al. 2018), millipedes (Loria et al. 2011), and beetles (Gomez et al. 2016; Leray et al. 2019). Surprisingly underrepresented in phylogenetic studies are trechine cave beetles (Coleoptera, Carabidae, Trechini), which account for 18% of all described troglobiotic diversity in the United States and Canada (Niemiller et al. 2019).

Cave trechines are prominent in many terrestrial subterranean habitats in Asia, Europe, and North America, and they are an ideal model system to study the evolution and biogeography of subterranean life (Faille et al. 2010, 2014; Ribera et al. 2010; Rizzo et al. 2013; Chen et al. 2021). They are small, predatory ground beetles (3–8 mm long) almost all of which lack eyes, are flightless, and are depigmented with long, slender bodies, elongated appendages, and sensory setae (‘aphaenopsian’; Barr 2004; Ober et al. 2022). North American cave trechine beetles include 162 taxa in six genera distributed primarily in the Appalachians valley and ridge (APP), Interior Low Plateau (ILP), and Ozarks (OZK) karst regions of central and eastern North America (Barr 2004; Ober et al. 2022. The genus *Pseudanophthalmus* is exceptionally diverse with 155 described species and subspecies occurring throughout the APP and ILP and is arranged into 26 morphologically-defined species groups (Barr 2004); however, there may be more than 80 additional undescribed species in this genus, including one species in the OZK (Peck 1998; Barr 2004; Ober et al 2022). Unlike *Pseudanophthalmus*, species richness in the other cave trechine genera is low, including *Ameroduvalius* (one species in the ILP), *Darlingtonea* (one species in the ILP), *Neaphaenops* (1 species in the ILP), *Nelsonites* (two species in the ILP), and *Xenotrechus* (two species in the eastern OZK). These five genera are thought to be closely related to *Pseudanophthalmus* (Valentine 1952; Barr 1972, 1980, 1981, 1985b; Maddison et al. 2019), but phylogenetic relationships among them remain unclear. Moreover, all genera but *Xenotrechus* co-occur with *Pseudanophthalmus*. All but 19 described cave trechines in North America are at an elevated risk of extinction (NatureServe 2022), in part because most species have exceptionally small ranges <10,000 km^2^ (i.e., short-range endemism sensu Harvey 2002), and many are known from just one or a few cave systems (Culver et al. 2000, 2003; Barr 2004; Niemiller and Zigler 2013; Niemiller et al. 2017; Malabad et al. 2021; Harden et al. 2022).

Several hypotheses have been proposed related to the diversification and biogeography of temperate North American cave fauna (Barr and Holsinger 1985; Holsinger 2000), including cave trechines (recently reviewed in Ober et al. 2022). These hypotheses largely group into two contrasting (but not mutually exclusive) scenarios. Speciation in cave organisms is traditionally thought to occur at the surface-cave ecotone when subterranean populations diverge from related surface populations (Barr 1985a; Barr and Holsinger 1985; Holsinger 2000). Due to limited dispersal ability or significant barriers to dispersal, multiple, closely related subterranean species are the product of several independent subterranean colonization events from one or more surface ancestors followed by isolation and speciation without significant subterranean dispersal (i.e., the multiple origins hypothesis). Speciation also could occur in subterranean habitats whereby a small number of subterranean colonization events by one or more surface ancestors followed by isolation and diversification but also with subsequent subterranean dispersal and diversification (i.e., few origins hypothesis). Several recent studies have suggested that this mode of speciation may be more common than previously thought. The discovery of monophyletic groups comprised of many subterranean lineages has been inferred as strong evidence for the role of subterranean speciation after a single or a few colonization events by surface ancestors (Holsinger 2000; Faille et al. 2010; Juan et al. 2010; Ribera et al. 2010). Subterranean speciation is generally thought to be the product of limited dispersal through subterranean corridors followed by isolation causing vicariance (Barr and Holsinger 1985; Holsinger 2000; Ribera et al. 2010). However, the evolutionary and biogeographic mechanisms factors that facilitate subterranean speciation are not well known or well-studied, as most investigations of speciation in cave organisms have focused on the morphological and evolutionary changes that accompany invasion and colonization from the surface (Holsinger 2000; Juan et al. 2010).

Determining the biogeographic and evolutionary history of a group of organisms is difficult when related sister lineages are either extinct or remain unsampled. For many groups of subterranean organisms, this is especially problematic as surface relatives are extinct, making distinguishing between single versus multiple colonization scenarios and distinguishing between modes of speciation (i.e., surface-subterranean ecotone versus subterranean vicariance) impossible using phylogenetic evidence alone. Monophyly of many subterranean taxa may reflect multiple, independent subterranean colonization events by a single surface ancestor with little to no subsequent subterranean dispersal and speciation or a single subterranean colonization event with substantial subterranean dispersal and speciation. Except for *P. sylvaticus*, which is known from deep soil of a high elevation spruce forest in West Virginia (Barr 1967b), no obvious surface-dwelling sister group is known in eastern North America for North American cave trechines.

*Pseudanophthalmus* and the other four troglobiotic genera belong to the *Trechoblemus* series in the *Trechus* assemblage (Jeannel 1926, 1927, 1928, 1930, 1949). *Trechoblemus* is predominately a Eurasian genus of surface-dwelling and winged trechine beetles with a single species *T. westcotti* Barr,1972 known from the Willamette Valley of Oregon (Barr 1972), which is closest known surface relative to North American cave trechines (Ober et al. 2022). Although there are few studies that discuss the genetic diversity and phylogeny of cave-dwelling carabid beetles in North America (Gómez et al. 2016; Boyd et al. 2020), a comprehensive study investigating the phylogeny, divergence time, and biogeography of cave carabid beetles is not yet available. The evolutionary and colonization history of subterranean habitats of eastern North American trechine beetles is complex with many questions remaining unanswered. More research is required to understand the diversification, origin, molecular systematics, and biogeographic patterns of cave trechines in eastern North America.

The aim of this study is to generate the first molecular phylogenetic framework for the study of the origin, diversification, and biogeography of cave trechines in eastern North America using ultraconserved elements (UCEs) phylogenomics. Cave beetle distributions and speciation likely have been influenced by both intrinsic (e.g., dispersal ability, body size, degree of subterranean adaptation) and extrinsic (e.g., vicariant events and habitat connectivity) factors (Juan et al. 2010; Porter 2007). Intrinsic factors may be more relevant at smaller spatial scales, such as within and among cave systems, and occurring over ecological timescales (Porter 2007), whereas extrinsic factors may be more relevant at larger spatial scales, such as within karst regions, and longer evolutionary timescales. Our objectives were to: (1) reconstruct the phylogenetic relationships of cave trechines using UCEs to examine monophyly and relationships of morphological species groups; (2) conduct divergence time analyses to determine relative timing of diversification particularly with respect to predictions of the climate-relict hypothesis; and (3) conduct biogeographic analyses to reconstruct colonization history and dispersal.

## Materials and methods

### Study area

Caves and subterranean fauna in the central and eastern United States are primarily associated with three major karst biogeographic regions (Culver and Hobbs 2002; Culver et al. 2003; Niemiller et al. 2019): the Appalachian Ridge and Valley, also called Appalachians (APP), the Interior Low Plateau (ILP), and the Ozarks (OZK) (Barr and Holsinger 1985; Culver et al. 2000; Figure 1). Rock strata in the ILP and OZK are mostly horizontally-bedded, whereas the rock layers in the APP were significantly faulted and folded due to past tectonic events associated with the uplift of the Appalachian Mountains. The ILP possesses the greatest number and density of caves, while the OZK is the largest in terms of surface area. The APP is ca. 37,000 km^2^ extending from southeastern New York to northeastern Alabama and includes a series of parallel sandstone ridges with intervening carbonate valleys. The ILP is ca. 61,000 km^2^ covering a large region west of the Cumberland Plateau from southern Indiana and Illinois southward through central Kentucky, central Tennessee, and northern Alabama (Ober et al. 2022). The OZK is ca. 129,500 km^2^ covering southern Missouri and northern Arkansas and features flat to gently fold cherty sandstone and cherty dolomite (McKenney and Jacobson 1996). The APP and ILP karst regions are proximate to one another, and they come in contact near the junction of the borders of Tennessee, Alabama, and Georgia. Both karst regions contain prominent karst exposures near the surface and support numerous caves (Culver et al. 2003; Weary and Doctor 2014). The ILP (400+ taxa) and APP (320+ taxa) possess the greatest troglobiotic species richness in the United States while OZK ranks fourth with 115+ species (Culver et al. 2003; Hobbs 2012; Niemiller et al. 2019).

**Figure 1.**
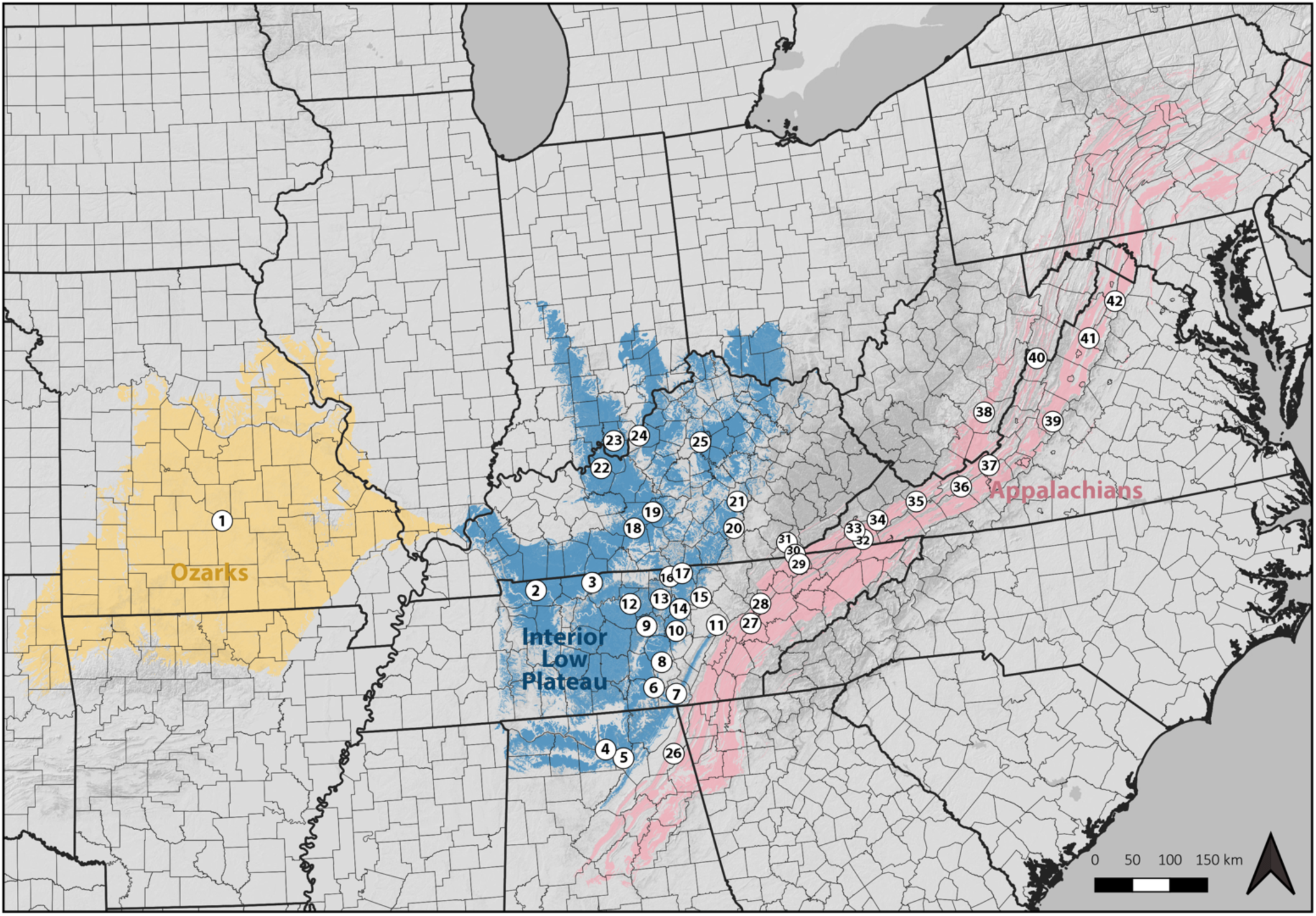
Map of the localities sampled for the studied species in the cave trechine group are indicated in white circles enclosing specimen number. Geographical area ranges including the Appalachians (APP), the Interior Low Plateau (ILP), and the Ozarks (OZK) are represented in pink, blue, and yellow colors respectively.

### Specimen sampling

We collected specimens of 45 cave trechine taxa from caves in the APP, ILP, and OZK, including 41 species of *Pseudanophthalmus* from 20 of the 26 species groups defined by Barr (2004), as well as *Ameroduvalius jeanneli*, *Darlingtonea kentuckensis*, *Neaphaenops tellkampfi*, *and Nelsonites jonesi* (Supplemental Table 1; Figure 1). We were not able to obtain specimens of *Nelsonites walteri* nor both species of *Xenotrechus*. Beetles were collected from terrestrial and riparian cave habitats, such as mud banks, the splash zones of active drips, near streams and rimstone pools, underneath rocks and coarse woody debris, or within cobble and gravel. Two species of the genus *Trechus — T. obtusus* and *T. humbolti —* and *Trechoblemus westcotti* were included as outgroups (Supplemental Table 1)

### DNA extraction, UCE library preparation, and sequencing

To generate a reduced representation genomic dataset of Ultraconserved Element (UCE) loci of cave trechines, we employed UCE phylogenomics (Faircloth et al. 2012; Branstetter et al. 2017), an approach that combines the targeted enrichment of thousands of nuclear UCE loci with multiplexed next-generation sequencing. Enrichment was performed using a published bait set that targets loci shared across all Coleoptera (‘Coleoptera 1.1Kv1’; Faircloth 2017). It includes 13,674 unique baits targeting 1,172 UCE loci. Whole genomic DNA was extracted from muscle, meso-metathorax, and leg of the cave trechine specimens following the DNA extraction method from Maddison et al. 1999. The extracted DNA samples were shipped to Arbor Biosciences (Ann Arbor, Michigan, USA) for downstream processing which included library preparation, target enrichment, and Illumina high throughput sequencing.

### Bioinformatics processing

The raw demultiplexed fastq reads was cleaned and processed using the software package Phyluce v1.7.1 (Faircloth 2016) and associated programs. We used Illumiprocessor v2.0 (Faircloth 2013), which is a parallel wrapper of Trimmomatic v0.40 (Bolger et al. 2014) to clean and trim raw reads. All programs hereafter beginning with ‘phyluce’ are python programs part of the Phyluce package After trimming, we generated summary stats on the trimmed reads using the Phyluce script ‘*phyluce_assembly_get_fastq_lengths*’. We assembled the cleaned/trimmed reads using ‘*phyluce_assembly_assemblo_abyss*’ with the AbySS assembler v2.3.5 (Simpson et al. 2009) using a kmer value of 60 on the CaveBio lab workstation. Next, we generated summary statistics (counts/lengths) of the assembled contigs using ‘*phyluce_assembly_get_fasta_lengths*’. To identify UCE loci, we used ‘*phyluce_assembly_match_contigs_to_probes*’ that incorporates Lastz v1.0 (Harris 2007) to match the Coleoptera v1 probe set sequences to contig sequences with a minimum coverage of 75% and minimum identity of 75% and created a relational database of hits. We used ‘*phyluce_assembly_get_match_counts*’ to create an initial database of loci counts per taxon. Next, we used ‘*phyluce_assembly_get_fastas_from_match_counts*’ to get a count of the number of UCE loci captured for each taxon. We then divided each UCE loci into a separate fasta file using ‘*phyluce_assembly_explode_get_fastas_file*’, as we later wanted to construct individual gene tree phylogenies, and then aligned the sequences and trimmed the edges using ‘*phyluce_align_seqcap_align*’ which implements MAFFT v7.130b (Katoh and Standley 2013). To remove poorly aligned regions, we trimmed internal and external regions of alignments using ‘*phyluce_align_get_gblocks_trimmed_alignments_from_untrimmed*’ that incorporates Gblocks (Talavera & Castresana 2007), with reduced stringency parameters (b1:0.5, b2:0.5, b3:12, b4:7). Next, we generated summary statistics on the alignments using ‘*phyluce_align_get_align_summary_data*’. Lastly, we used ‘*phyluce_align_get_only_loci_with_min_taxa*’ and ‘*phyluce_align_concatenate_alignments*’ to filter and create two concatenated data sets or matrix by selecting aligned loci that contains at least 50% and 75% of the total taxa for phylogenetic inference.

### UCE phylogenomics

We performed two different types of phylogenomic analyses: (1) concatenated analyses using RaxML-HPC2 Workflow on XSEDE v8.2.12 (Stamatakis 2014) on the CIPRES Science Gateway v3.3 (https://www.phylo.org/) and MrBayes v3.2 (Ronquist et al. 2012) module of PhyloSuite v1.1.15 (Zhang et al. 2019) (2) species tree analyses using ASTRAL-II v5.7.8 (Mirarab and Warnow 2015) and SVDQuartets (Chifman and Kubatko 2014) implemented in PAUP* v4.0a169 (Swofford 1998). For all subsequent analyses, we used the 75% complete data-matrix.

For concatenated analyses, we defined each UCE locus as its own character set and then used PartitionFinder v2.1.1 (Lanfear et al. 2017) implementing the ‘greedy’ search algorithm (Lanfear et al. 2012) to select for the best partitioning strategy for the data under the General Time Reversible + Gamma (GTRGAMMA) site rate substitution model using the AICc metric (Burnham and Anderson 2002). We then conducted 20 maximum-likelihood (ML) searches in RaxML-HPC2 Workflow on XSEDE (Towns et al. 2014). We also performed non-parametric bootstrap replicates under GTRGAMMA using the autoMRE option to optimize the number of bootstrap replicates for this large dataset. We reconciled the bootstrap replicates with the best fitting ML tree. To confirm the reliability of the tree topology, the concatenated dataset was also processed through MrBayes for Bayesian phylogenetic reconstruction. We used the same partitioning strategy described above and estimated the most appropriate site rate substitution model for MrBayes using PartitionFinder v2.1.1. We conducted two independent runs of one cold chain and three heated chains (default settings) for 5,000,000 Markov chain Monte Carlo (MCMC) generations sampling every 100 generations in MrBayes. After dropping the first 25% ‘burn-in’ trees to ensure stationarity and examining the log-likelihood values for each Bayesian run using Tracer v1.7 (Rambaut et al. 2018), the remaining 37,500 sampled trees were used to estimate the consensus tree and the associated Bayesian posterior probabilities. A midpoint rooted ML and Bayesian trees with support values were generated using FigTree v1.4.4 (Rambaut 2010). To evaluate if allowing missing taxa on total alignment length affected the topology of the tree, the above-mentioned analyses were also conducted on the 50% complete data-matrix.

For species tree analyses, we first reconstructed the ML gene tree estimated for each of the UCE loci in IQTree v1.6.8 (Nguyen et al. 2015) module of PhyloSuite under the General Time Reversible + Gamma (GTRGAMMA) site rate substitution model. We conducted 1,000 iterations as well as 5,000 non-parametric ultrafast bootstrap replicates. We reconciled the bootstrap replicates with the best fitting ML tree of each locus. Second, these gene trees were input to ASTRAL-II to create a multispecies coalescent species tree and assessed support with 150 bootstrap replicates creating support values akin to posterior probabilities of nodes. Finally, we used another species tree method to look for congruence between methods. We created a multispecies coalescent species tree using SVDQuartets in PAUP* where we evaluated 100,000 random quartets using the Quartet FM (QFM) algorithm (Reaz et al. 2014). We assessed support with 200 bootstrap replicates. A midpoint rooted ASTRAL and SVDQuartets species trees with support values were generated using FigTree v.1.4.4.

### Estimation of divergence times

To estimate the relative age of divergence of the cave trechine lineages, we used the Bayesian relaxed phylogenetic approach implemented in BEAST2 v2.6.7 (Drummond and Rambaut 2007), which allows for variation in substitution rates among branches (Drummond et al. 2006). We implemented a GTR+G model of DNA substitution with four rate categories for all partitions of the 75% complete data-matrix. We chose the uncorrelated lognormal relaxed molecular clock model to estimate the substitution rates and the Yule process of speciation as the tree prior. Because of the absence of a robust closely related fossil records for trechine beetles, we used a secondary calibration method to date the trees. Thus, we set the ‘ucldMean’ prior a lognormal distribution with average equal to a per-branch rate of 0.0012 substitutions/site/MY and a standard deviation of 0.059. All other parameters including the ‘ucldStdev’ prior was left with default settings. This rate is obtained from Faille et al. (2013) for the subterranean Trechini based on the colonization of the Alps in Europe (0.0010 and 0.0013 substitutions/site/MY for the nuclear small ribosomal unit 18S rRNA and large ribosomal unit 28S rRNA, respectively).

We performed two independent runs for 25,000,000 generations sampling every 500 generations in BEAST2. For both runs, we assessed convergence, likelihood, stationarity and verified effective sample size (ESS) values of each parameter using Tracer v1.7. After discarding a burn-in of 10%, results of both runs were combined with LogCombiner v2.6.7 and the consensus tree was compiled with TreeAnnotator v2.6.7 (Drummond and Rambaut 2007). A midpoint rooted tree with geological time scale depicting the node ages, 95% highest posterior density interval (HPD) bars, and posterior probabilities was generated using the R strap package (R Development Core Team, 2014) using a custom R script.

### Ancestral range estimation

We estimated ancestral ranges using stochastic likelihood-based models of geographic range evolution implemented in the R package ‘BioGeoBEARS 0.2.1’ (Matzke 2013). We executed and compared the standard dispersal-extinction-cladogenesis (DEC) model with the (DEC+J) model (Ree et al. 2005; Ree and Smith 2008), which includes an additional parameter j that allows for founder-event speciation by jump dispersal (Matzke 2014). The ‘j’ parameter allows for a daughter lineage to immediately occupy via long-distance dispersal a new area that is different from the parental lineage. We used the maximum clade credibility time-calibrated tree from the concatenated BEAST2 analysis as the input tree. After pruning the outgroup species, we assigned each species in the tree the biogeographic major karst regions (represented by single letter code) spanned by the represented lineage: Appalachian Ridge and Valley (A), the Interior Low Plateau (I), and the Ozark Highlands (O). In addition, to have a closer look at the effect of different karst subregions on the represented lineage, we conducted an additional analysis by assigning each species in the tree to biogeographic karst subregions: Ridge and Valley (R), Wills Valley (V), Greenbrier Karst (G), Inner Bluegrass (I), Western Pennyroyal (W), Highland Rim (H), Pine Mountain (P), Cumberland Plateau (C), Nashville Basin (N), Outer Bluegrass (O), Sequatchie Valley (S), and Ozarks (Z). Each species was coded as being present or absent in each of these areas and the maximum number of areas occupied by a single species was set to 4. The models were compared to each other using two different methods: (1) the likelihood of each model was compared with Akaike Information Criterion (AIC); and (2) the nested models were compared with each other using a chi-squared test to determine if the model with ‘j’ parameter was preferred. Biogeographic stochastic mapping (Dupin et al. 2017) using the best-fitting biogeographic model (DEC+J) was plotted on the cave trechine chronogram, and the number of dispersal events among different karst regions was assessed.

## Results

Information on specimen vouchers, DNA quantities, raw Illumina reads before and after quality filtering and trimming, and SRA accession numbers can be found in Supplemental Table 1. After filtering and trimming, our aligned UCE contigs included 3,240,300 characters, of which 2,885,448 were nucleotides and 354,852 (11.0%) were missing data. Mean locus length was 224 nucleotides (range: 42–798 bp). Each locus contained an average of 12 informative characters (range: 0–95). We analyzed both 50% and 75% coverage data matrices. The 75% UCE data matrix comprised 16,794 base pairs (bp) and 68 UCE loci for 48 specimens (45 cave trechine beetles and three outgroup taxa). The 50% UCE data matrix comprised 65,376 base pairs (bp) and 274 loci for the 48 specimens.

Raw Illumina sequence reads are available at the NCBI Short-Read Archive (BioProject PRJNA894729; https://www.ncbi.nlm.nih.gov/bioproject/PRJNA894729) while aligned data matrices and tree files are available at Dryad (doi:10.5061/dryad.sj3tx967w).

### Concatenated phylogenetic analyses

Both ML and Bayesian inference (BI) trees inferred from the 75% concatenated UCE data matrix resolved similar topologies with high support for most nodes (Figures 2 & 3). Likewise, ML and Bayesian inferred from the 50% concatenated UCE data matrix (Supplemental Figures 1 & 2) were highly congruent with trees inferred from the 75% concatenated UCE data matrix; therefore, we focus our discussion based on the results of concatenated (and species tree analyses in the next section) of the 75% UCE data matrix. Of the 15 *Pseudanophthalmus* species groups as defined by Barr (2004) for which more than one taxon was sampled, 12 were monophyletic. Four species groups were paraphyletic, including the *cumberlandus*, *grandis*, *jonesi*, and *simplex* species groups. Moreover, the genus *Pseudanophthalmus* was not recovered as monophyletic as all of the other cave trechine genera included in the analyses (i.e., *Ameroduvialis*, *Darlingtonea*, *Neaphaenops*, and *Nelsonites*) were nested within *Pseudanophthalmus*.

**Figure 2.**
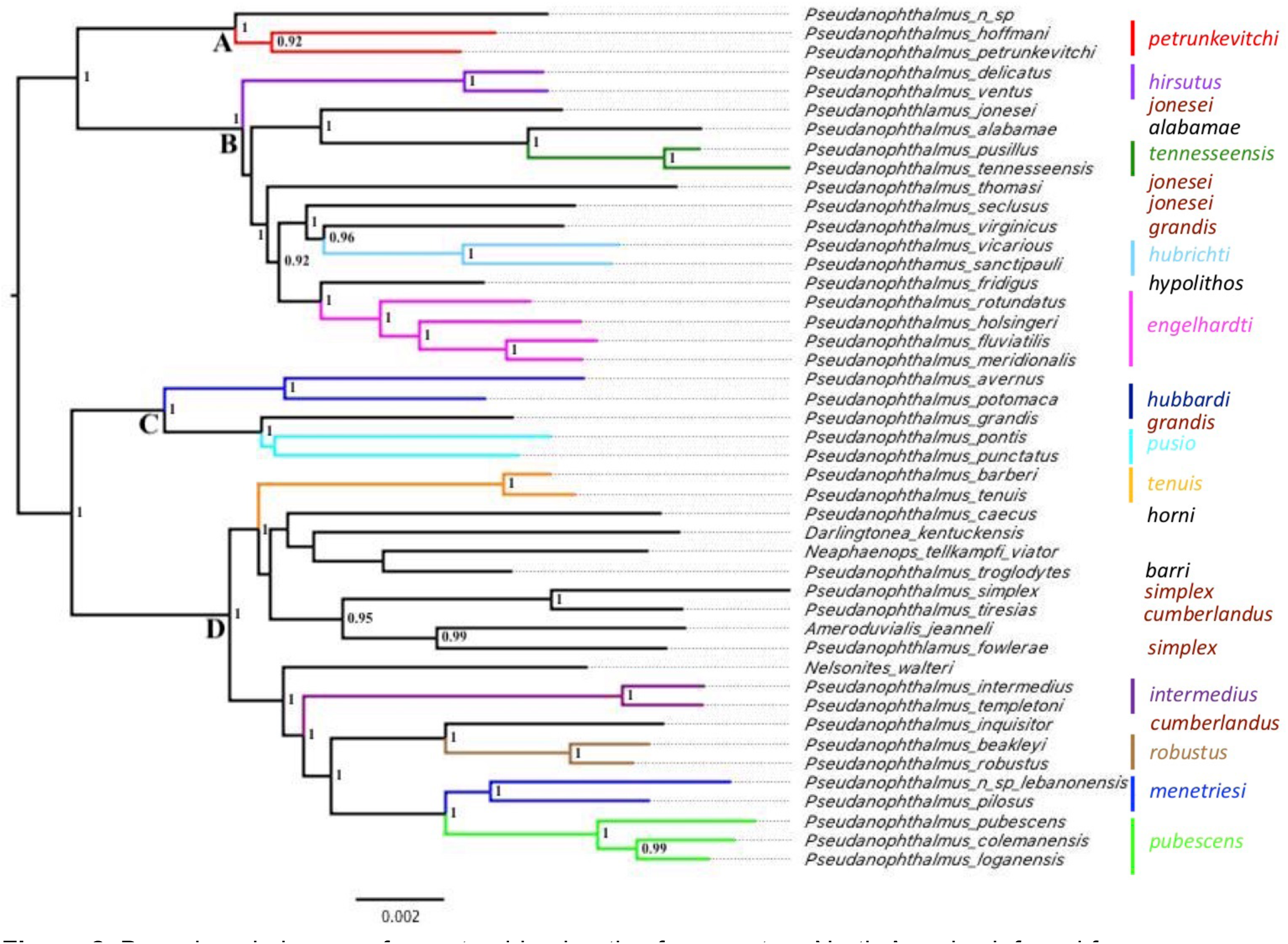
Bayesian phylogeny of cave trechine beetles from eastern North America inferred from 75% complete concatenated UCE matrix. Numbers indicate support values (Bayesian posterior probability) for nodes greater than 0.90. Outgroup taxa not shown, and four primary clades are labeled A, B, C, and D.

**Figure 3.**
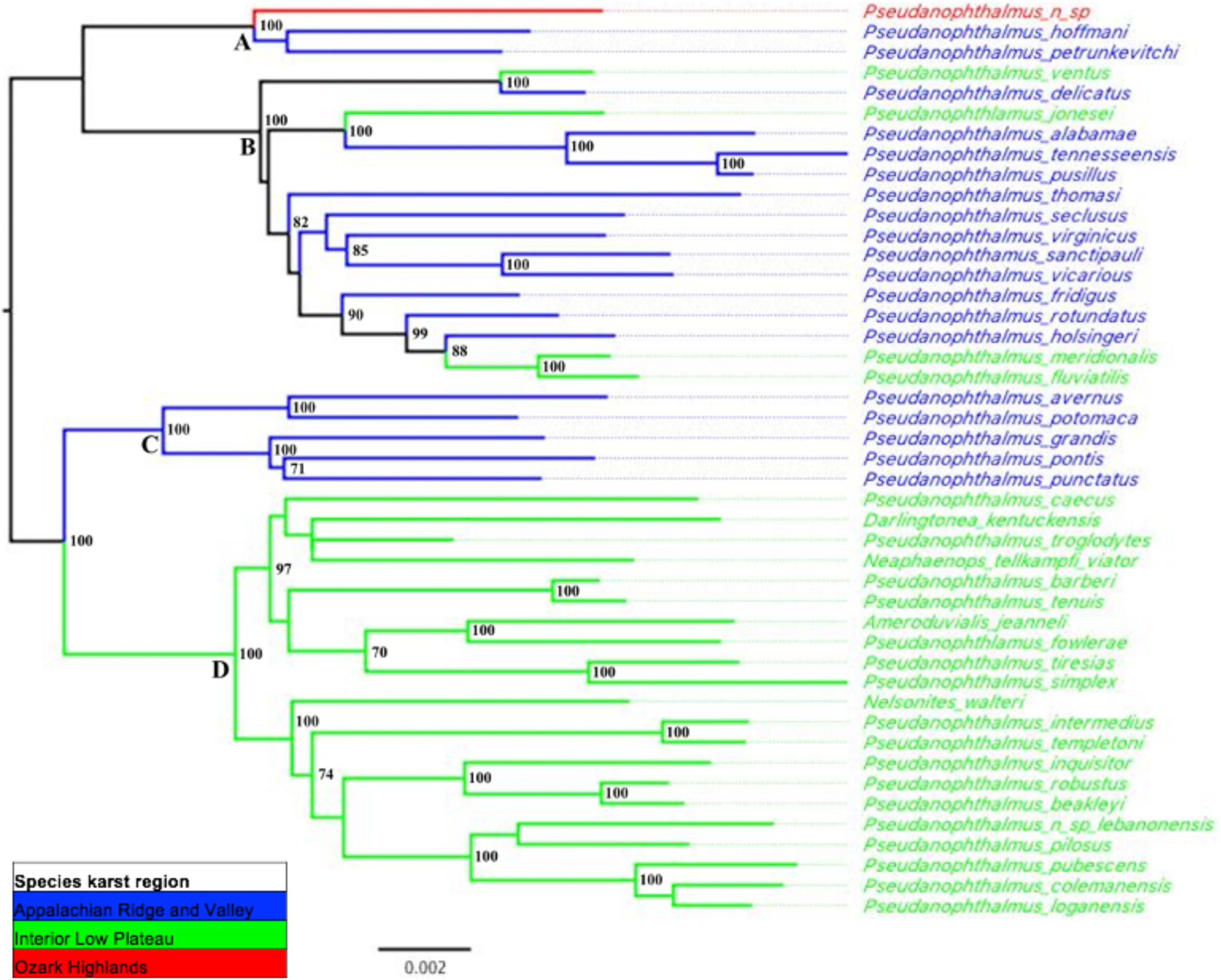
Maximum-likelihood phylogeny of cave trechine beetles from eastern North America inferred from 75% complete concatenated UCE matrix. Numbers indicate bootstrap support values (maximum-likelihood bootstrap) for nodes greater than 70. Outgroup taxa not shown, and four primary clades are labeled A, B, C, D.

Four primary clades were recovered in both ML and BI analyses (Clades A, B, C, and D in Figures 2 & 3, Supplemental Figures 1 & 2). *Pseudanophthalmus* formed two main clades that largely corresponds with the karst biogeographic regions: Appalachian Ridge and Valley and Interior Low Plateau. The genera *Ameroduvalius*, *Darlingtonea*, *Neaphaenops*, and *Nelsonites* were nested within the Interior Low Plateau *Pseudanophthalmus* clade. We found strong support for a clade with the geographically isolated undescribed *Pseudanophthalmus* species sister to the *P*.*petrunkevitchi* species group of the APP (Clade A). Clade B includes seven *Pseudanophthalmus* species groups (*engelhardti*, *hubrichti*, *tennesseensis, hirsutus, jonesi, alabamae,* and *hypolithos*) along with one species from the *grandis* species group (*P. virginicus*). The *P. hubbardi* and *pusio* species groups along with the other species from the *grandis* species group (*P. grandis*), all from the APP, form Clade C. Finally, Clade D is comprised of taxa from the ILP, including nine *Pseudanophthalmus* species groups (*tenuis, intermedius, robustus, menetriesi, pubescens, horni, barri, simplex,* and *cumberlandus*) as well as the genera *Ameroduvalius*, *Darlingtonea*, *Neaphaenops*, and *Nelsonites*. The weakly supported sister group relationships of the *P. barberi* and *P. tenuis* clade within Clade D was the only difference between the BI and the ML phylogenetic trees.

### Species tree analyses

The species tree inferred from the 75% UCE data matrix using ASTRAL-II differed slightly from concatenated ML and BI trees. Differences were not strongly supported, however, and generally involved the placement of *P. thomasi*, *P. fridigus*, *N. telkampfi viator*, *P. troglodytes*, and *D. kentuckensis* within primary clades (Figure 4). In general, support for several deeper nodes in the ASTRAL species tree was relatively low. The species tree reconstructed using SVDQuartets (Supplemental Figure 3) was similar in topology to the concatenated ML and BI trees (Figures 2 & 3). However, several deeper nodes in the SVDQuartets species tree were weakly supported.

**Figure 4.**
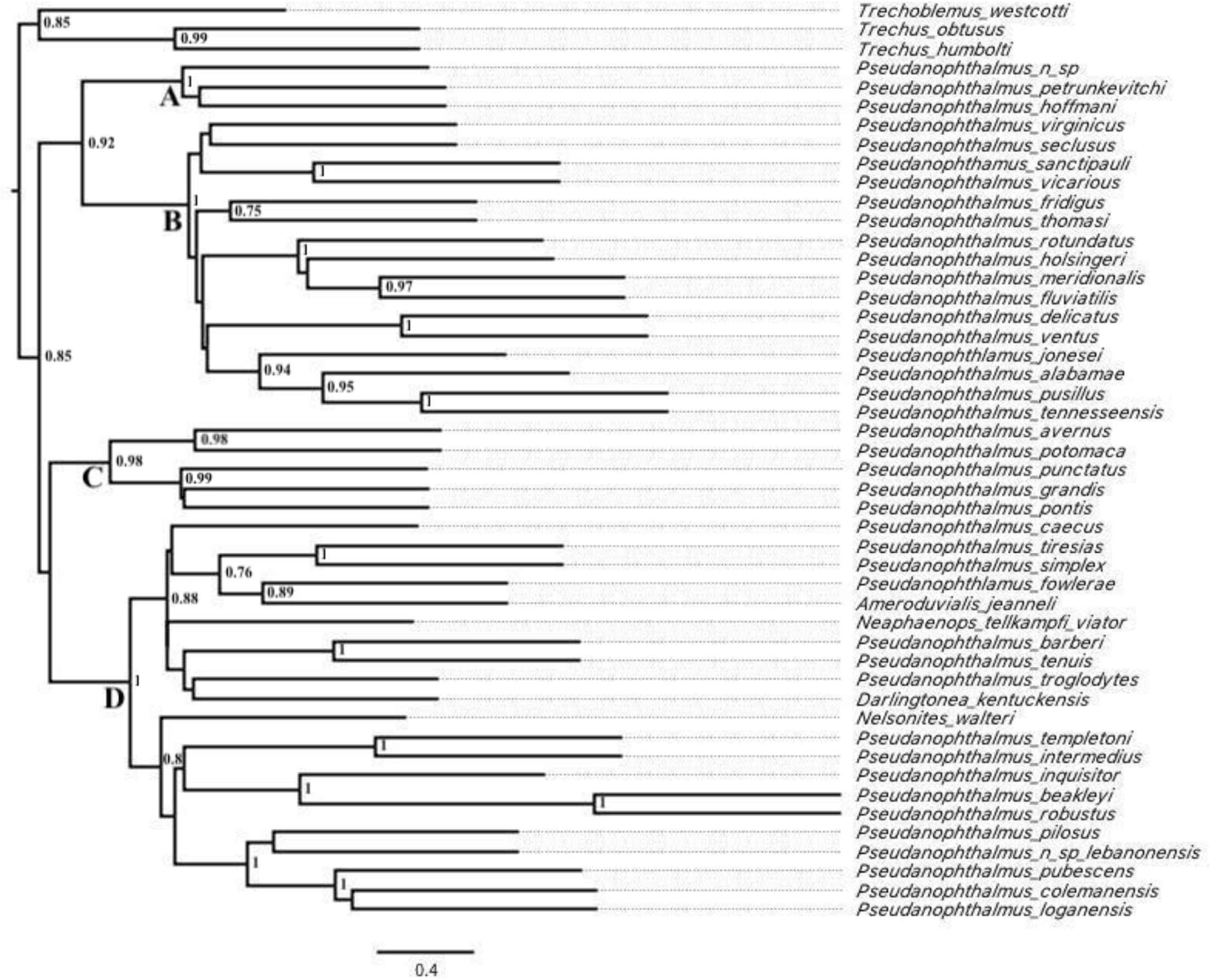
ASTRAL coalescent species tree, input trees derived from multi-partitioned IQTree analyses of individual gene trees. Lower posterior probability support values greater than 0.90 are displayed.

### Divergence time estimation

Divergence time estimation in BEAST2 using a molecular clock rate recovered a crown age for North American trechine beetles of 11.5 Mya (95% highest posterior density [HPD] interval: 9.8– 13.2 Mya) in the middle Miocene at the beginning of the Tortonian (Figure 5 and Supplemental Figure 4) forming the main ARV and ILP clades. Additional diversification of primary clades with the ARV and ILP clades occurred shortly thereafter at 9.5 Mya (95% HPD: 7.7–11.4 Mya) and 10.7 Mya (95% HPD: 9.1–12.4 Mya), respectively. Diversification within the primary clades occurred primarily from the late Miocene into the Pliocene with estimated species divergences all slightly predated the Pliocene (i.e., 5 Mya). The divergence events estimated to have occurred in the Pleistocene were limited primarily to species pairs within species groups (Figure 5 & Supplemental Figure 4). Diversification during the Pliocene and Pleistocene were almost exclusively within the same geographical area and involved species of the same morphological species group.

**Figure 5.**
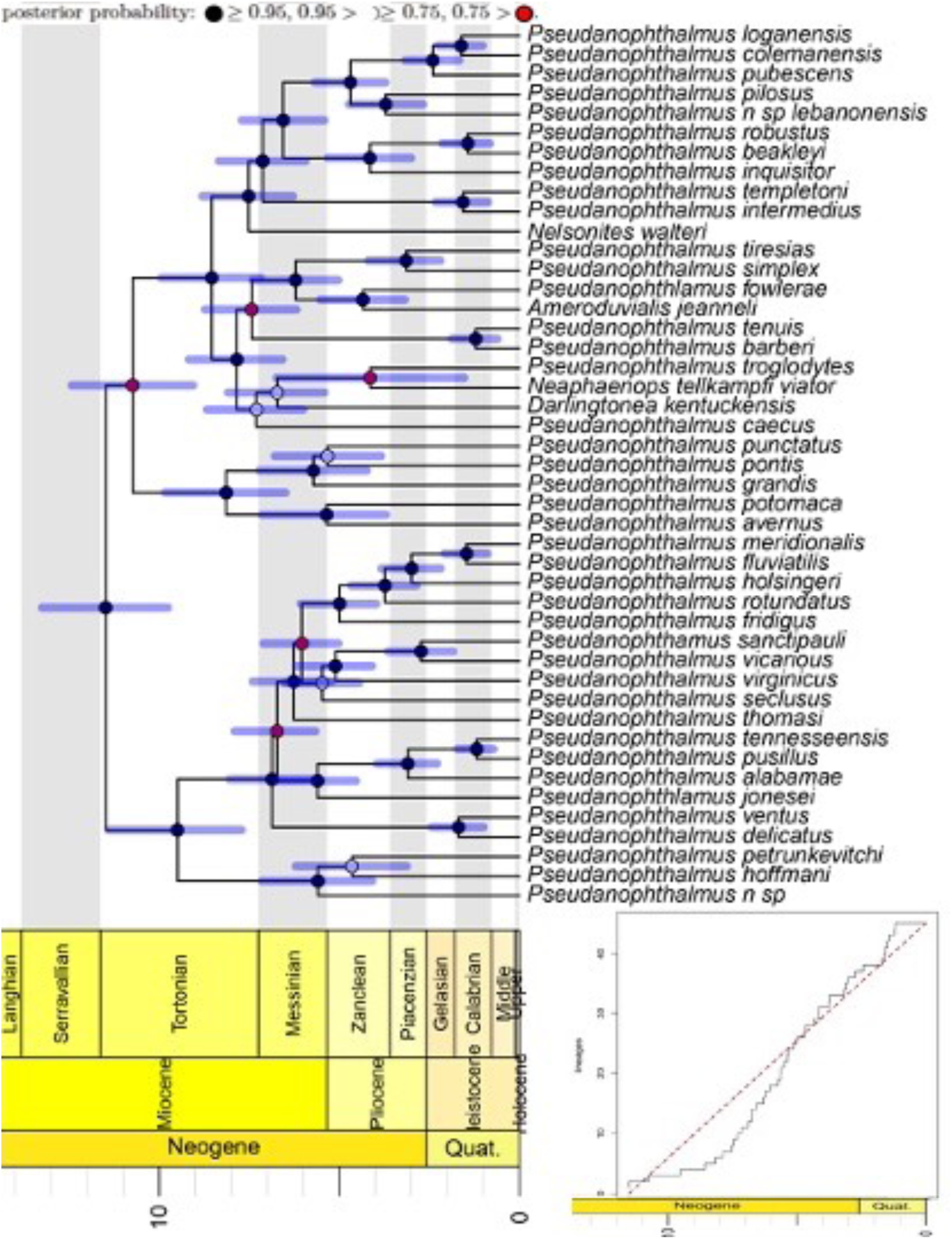
Time-calibrated maximum clade credibility tree inferred from 75% concatenated UCE matrix, summarized by TreeAnnotator, and plotted with a geological time scale using the strap package in R. Phylogeny dated using a Bayesian relaxed clock method in BEAST. Branches are proportional to time in millions of years. Outgroups were pruned after analyses for an enlarged view. The 95% confidence intervals for the ages of basal branches in the tree are indicated with transparent blue bars. Lineages through time plot displayed at lower right corner. The internal nodes of the tree are indicated with circles, where circles mark nodes with posterior probability:

### Ancestral range reconstruction

The DEC+J model provided a significantly better fit for the phylogeny resolved using BEAST compared to other karst region models (Table 2). The earliest divergence in cave trechine beetles was estimated to occur ∼11.5 Mya in a widespread common ancestor most likely occurring in Appalachian Ridge and Valley with dispersal into the Interior Low Plateau between 9.0–12.5 Mya (Figure 5). At least three additional colonization events from the Appalachians Ridge and Valley into the Interior Low Plateau are supported and are estimated to have occurred 2.0–5.5 Mya. The model also supports a single dispersal event from the Appalachian Ridge and Valley into the Ozark Highlands (Figure 6).

**Table 1.** Specimens used in the study, with species group information, locality including karst regions, and voucher reference numbers

| Specimen | Voucher | Species | Species-group | State | County | Cave | Karst_region | Karst_subregion |
| --- | --- | --- | --- | --- | --- | --- | --- | --- |
| 1 | DNA689 | *Pseudanopthalmus n.sp.* | ? | MO | Texas | Pine Hollow Cave | OZK | Ozarks |
| 2 | DNA370 | *Pseudanopthalmus colemanensis* | *pubescens* | TN | Montgomery | Clarksville Lake Cave | ILP | Western Pennyroyal |
| 3 | DNA498 | *Pseudanopthalmus loganensis* | *pubescens* | TN | Robertson | Bradley Hill Caverns | ILP | Western Pennyroyal |
| 4 | DNA385 | *Pseudanopthalmus fluviatilis* | *engelhardti* | AL | Morgan | Talucah Cave | ILP | Cumberland Plateau |
| 5 | DNA303 | *Pseudanopthalmus meridionalis* | *engelhardti* | AL | Marshall | Beech Spring Cave | ILP | Cumberland Plateau |
| 6 | DNA474 | *Pseudanopthalmus intermedius* | *intermedius* | TN | Franklin | Dry Cave | ILP | Cumberland Plateau |
| 7 | DNA570 | *Pseudanopthalmus ventus* | *hirsutus* | TN | Marion | Dancing Fern Cave | ILP | Sequatchie Valley |
| 8 | DNA571 | *Pseudanopthalmus templetoni* | *intermedius* | TN | Grundy | Skull Cave | ILP | Cumberland Plateau |
| 9 | DNA510 | *Pseudanopthalmus tiresias* | *cumberlandus* | TN | DeKalb | Indian Grave Point Cave | ILP | Highland Rim |
| 10 | DNA404 | *Pseudanopthlamus jonesei* | *jonesei* | TN | Cumberland | McCullough Sump Cave | ILP | Sequatchie Valley |
| 11 | DNA315 | *Pseudanopthalmus robustus* | *robustus* | TN | White | John Henry Demps Cave | ILP | Cumberland Plateau |
| 12 | DNA289 | *Pseudanopthalmus n. sp. lebanonensis* | *menetriesi* | TN | Wilson | Shell Caverns | ILP | Nashville Basin |
| 13 | DNA600 | *Pseudanopthalmus simplex* | *simplex* | TN | Jackson | Flatt Cave | ILP | Highland Rim |
| 14 | DNA240 | *Nelsonites walteri* |  | TN | Putnam | Stamps Cave | ILP | Cumberland Plateau |
| 15 | DNA511 | *Pseudanopthalmus beakleyi* | *robustus* | TN | Fentress | Hurricane Maze Cave | ILP | Cumberland Plateau |
| 16 | DNA410 | *Pseudanopthlamus fowlerae* | *simplex* | TN | Clay | Shankey Branch Cave | ILP | Highland Rim |
| 17 | DNA574 | *Pseudanopthalmus inquisitor* | *cumberlandus* | TN | Clay | JC Melton Cave | ILP | Cumberland Plateau |
| 18 | DNA257 | *Pseudanopthalmus pubescens* | *pubescens* | KY | Barren | L & N Railroad Cave | ILP | Western Pennyroyal |
| 19 | DNA656 | *Neaphaenops tellkampfi viator* |  | KY | Green | Scotts Cave | ILP | Western Pennyroyal |
| 20 | DNA296 | *Ameroduvialis jeanneli* |  | KY | Pulaski | Drowned Rat Cave | ILP | Cumberland Plateau |
| 21 | DNA467 | *Darlingtonia kentuckensis* |  | KY | Rockcastle | Fletcher Spring Cave | ILP | Cumberland Plateau |
| 22 | DNA618 | *Pseudanopthalmus pilosus* | *menetriesi* | KY | Hardin | Cassell Cave | ILP | Western Pennyroyal |
| 22 | DNA607 | *Pseudanopthalmus barberi* | *tenuis* | KY | Breckinridge | Webster Cave | ILP | Western Pennyroyal |
| 23 | DNA525 | *Pseudanopthalmus tenuis* | *tenuis* | IN | Harrison | Binkley Cave | ILP | Western Pennyroyal |
| 24 | DNA544 | *Pseudanopthalmus troglodytes* | *barri* | KY | Jefferson | Eleven Jones Cave | ILP | Outer Bluegrass |
| 25 | DNA554 | *Pseudanopthalmus caecus* | *horni* | KY | Woodford | Richardson Spring Cave | ILP | Inner Bluegrass |
| 26 | DNA748 | *Pseudanopthalmus alabamiae* | *alabamiae* | AL | DeKalb | Manitou Cave | APP | Wills Valley |
| 27 | DNA313 | *Pseudanopthalmus tennesseensis* | *tennesseensis* | TN | Roane | Eblen Cave | APP | Ridge and Valley |
| 28 | DNA530 | *Pseudanopthalmus pusillus* | *tennesseensis* | TN | Anderson | Martin Cave | APP | Ridge and Valley |
| 29 | DNA593 | *Pseudanopthalmus rotundatus* | *engelhardti* | TN | Claiborne | Kings Saltpeter Cave | APP | Ridge and Valley |
| 29 | DNA492 | *Pseudanopthalmus delicatus* | *hirsutus* | TN | Claiborne | Kings Saltpeter Cave | APP | Ridge and Valley |
| 30 | DNA631 | *Pseudanopthalmus holsingeri* | *engelhardti* | VA | Lee | Young-Fugate Cave | APP | Ridge and Valley |
| 31 | DNA541 | *Pseudanopthalmus fridigus* | *hypolithos* | KY | Bell | Ice Box Cave | APP | Pine Mountain |
| 32 | DNA632 | *Pseudanopthalmus thomasi* | *jonesei* | VA | Scott | Blair-Collins Cave | APP | Ridge and Valley |
| 33 | DNA670 | *Pseudanopthalmus seclusus* | *jonesei* | VA | Scott | Kerns No.1 Cave | APP | Ridge and Valley |
| 34 | DNA758 | *Pseudanopthalamus sanctipauli* | *hubrichti* | VA | Russell | Banners Corner Cave | APP | Ridge and Valley |
| 35 | DNA685 | *Pseudanopthalmus virginicus* | *grandis* | VA | Tazewell | Hugh Young Cave | APP | Ridge and Valley |
| 35 | DNA766 | *Pseudanopthalmus vicarious* | *hubrichti* | VA | Tazewell | Ward Cave | APP | Ridge and Valley |
| 36 | DNA781 | *Pseudanopthalmus hoffmani* | *petrunkevitchi* | VA | Bland | Banes Spring Cave | APP | Ridge and Valley |
| 37 | DNA582 | *Pseudanopthalmus punctatus* | *pusio* | VA | Giles | Smokehole Cave | APP | Ridge and Valley |
| 38 | DNA538 | *Pseudanopthalmus grandis* | *grandis* | WV | Greenbrier | Culverson Creek Cave | APP | Greenbrier Karst |
| 39 | DNA577 | *Pseudanopthalmus pontis* | *pusio* | VA | Rockbridge | Bradys Cave | APP | Ridge and Valley |
| 40 | DNA756 | *Pseudanopthalmus potomaca* | *hubbardi* | VA | Highland | Vandevander Cave | APP | Ridge and Valley |
| 41 | DNA507 | *Pseudanopthalmus avernus* | *hubbardi* | VA | Rockingham | Endless Caverns | APP | Ridge and Valley |
| 42 | DNA767 | *Pseudanopthalmus petrunkevitchi* | *petrunkevitchi* | VA | Warren | Brother Daves Cave | APP | Ridge and Valley |
|  | DNA787 | *Trechus obtusus* | *outgroup* | OR |  |  |  |  |
|  | DNA786 | *Trechoblemus westcotti* | *outgroup* | OR |  |  |  |  |
|  | DNA788 | *Trechus humboldti* | *outgroup* | OR |  |  |  |  |

**Table 2.** Comparison of dispersal-extinction-cladogenesis (DEC) models with jump dispersal (+J) and without (+J) for cave trechine beetles based on their dispersal within major karst region. Abbreviations as follows: LnL, loglikelihood; numparams, number of parameters in each model; d, dispersal rate; e, extinction rate; j, founder-event speciation rate; AIC, Akaike Information Criterion; LRT, likelihood-ratio test.

| Model | LnL | numparams | d | e | j | AIC | AIC_wt | LRT<br>pval |
| --- | --- | --- | --- | --- | --- | --- | --- | --- |
| DEC | -30.31 | 2 | 0.0085 | 1.00E-12 | 0 | 64.62 | 0.0003 |  |
| DEC+J | -21.31 | 3 | 1.00E-12 | 1.00E-12 | 0.03 | 48.63 | 1 | 2.20E-05 |

**Figure 6.**
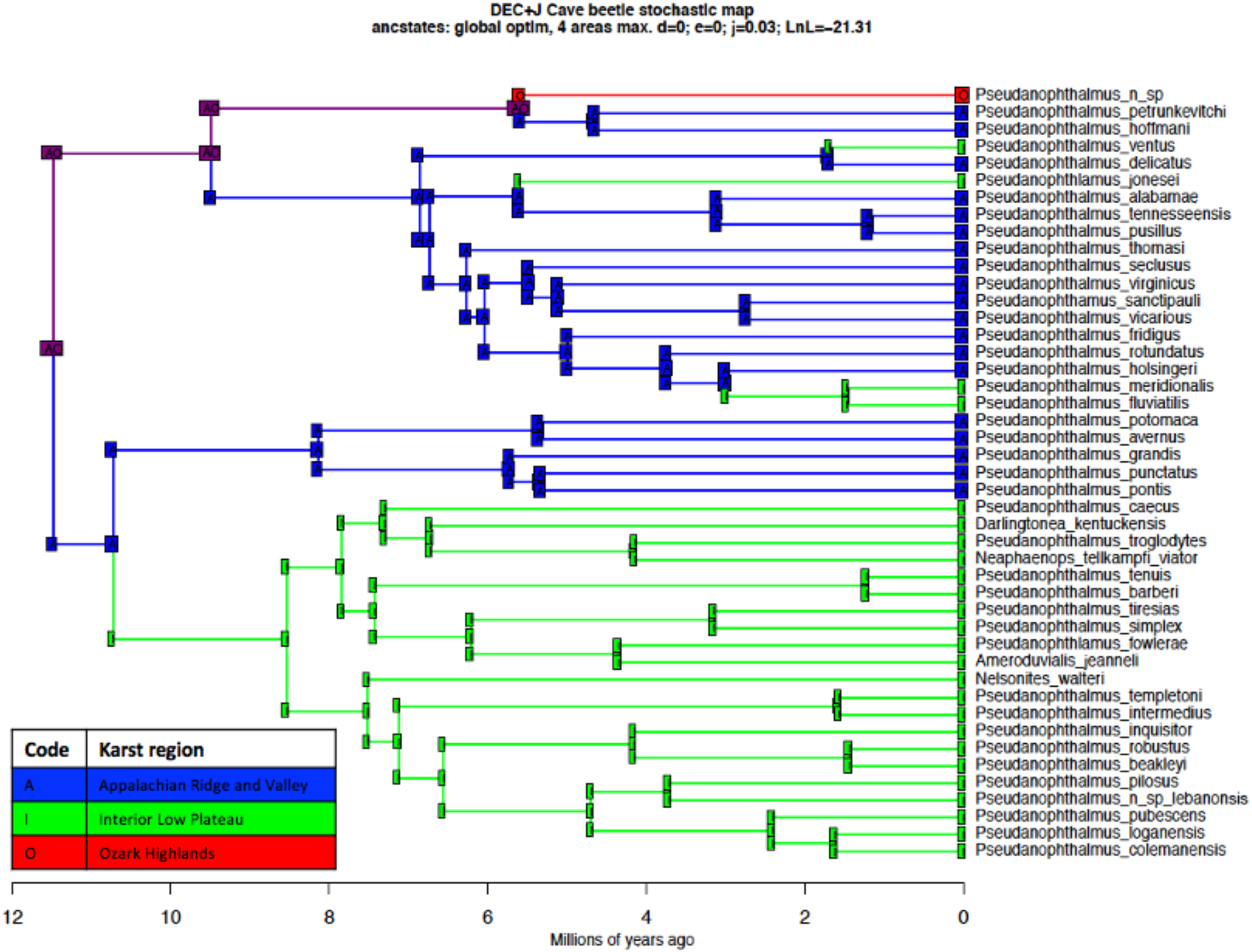
Ancestral area estimation for cave trechine beetles from eastern North America based on the preferred DEC+J model. Ancestral areas were estimated across the time-calibrated phylogeny inferred from 75% complete concatenated UCE matrix. Most probable ancestral karst region range at each node shown. Corner positions represent the geographic karst region range immediately after a cladogenetic event.

With respect to karst subregions, the DEC + J model again was the best model (AIC = 151.9; Supplemental Table 1). This model also supported a common ancestor in the Appalachian Ridge and Valley 11.5 Mya (95% HPD: 9.5–13.5 Mya) during the middle Miocene (Figure 6). The *petrunkevitchi* species group was the first group to colonize the Appalachian Ridge and valley karst and its sister lineage later dispersed into the Ozark Highlands. The main radiation in the Interior Low Plateau appears to have occurred from east to west first into the Cumberland Plateau then into more western ILP karst subregions, including the Highland Rim, Western Pennyroyal, Nashville Basin, Inner Bluegrass, and Outer Bluegrass. The general pattern of diversification within the Ridge and Valley clade suggests dispersal from north to south with occasional dispersal into adjacent ILP karst subregions, including the Cumberland Plateau, Pine Mountain, and Sequatchie Valley.

## Discussion

This study provided the first phylogenomic framework for the North American cave trechine beetles improving upon previous studies with expansive taxon and data sampling, resulting in a robust hypothesis of evolutionary relationships. Our results based on phylogenetic analyses of 68 genomic UCE loci shed new light on the systematics of carabid beetles and highlight potential areas for future taxonomic research. The study also presents a comprehensive analysis of biogeographic history for these unique beetles for the first time.

### Carabidae UCEs

Concatenated analyses in this study yielded more well-resolved trees than coalescent-based species tree methods (i.e., ASTRAL and SVDQuartets). Lower support values for some deeper branches in coalescent-based species tree approaches may be due to gene-tree discordance caused by incomplete lineage sorting (Maddison 1997; Edwards 2009), lack of phylogenetic signal among gene trees, and missing data (Thomson et al., 2008; Edwards 2009; Gatesy and Springer 2014; Springer and Gatesy 2014, 2016; Xi et al. 2015; Edwards et al. 2016; Meiklejohn et al. 2016; Moyle et al. 2016). Previous studies have demonstrated that species tree methods including ASTRAL and SVDQuartets can be prone to errors in gene tree estimation (Edwards et al. 2016; Hosner et al. 2016; Meiklejohn et al. 2016; Springer and Gatesy 2016). Hosner et al. (2016) showed that missing data can also cause significant errors in species tree estimation, especially when including taxa with only partially captured UCE loci. The length of a given UCE locus obtained via targeted capture methods may vary greatly among samples (Hosner et al. 2016; Streicher et al. 2016). In our dataset, the 50% and 75% complete dataset contained considerably fewer UCEs than other published beetle UCE datasets (Baca et al. 2017; Van Dam et al. 2017; Gustafson et al. 2020; Bradford et al. 2022; Sota et al. 2022) suggesting the potential for the missing data to affect our coalescent analyses. Other several studies have shown that there is a positive trade-off in constructing a larger data matrix by allowing the inclusion of loci with missing taxa when conducting concatenated analyses (Hosner et al. 2016; Streicher et al. 2016).

Therefore, the amount of missing data in our analyses may possibly account for the differences in resolution between our concatenated and coalescent analyses. No genomes for carabid beetles were publicly available for use during the development of the Coleoptera bait set (Faircloth 2017), which may account for the lower recovery of UCEs in our study.

### Evolutionary history and biogeography of North American cave trechine beetles

Distinguishing between the two hypotheses (multiple vs. few origins) and the continuum of colonization, dispersal, and speciation scenarios in between is not trivial and often requires other sources of data, such as geological, climatical, or paleontological evidence, to reconstruct phylogeographic histories of cave organisms. Evidence in support of either scenario in North American cave trechines from past studies is limited. Barr (2004) proposed that varying levels of troglomorphy observed among North American cave trechine beetles were indicative of multiple episodes of cave colonization, with the assumption that degree of troglomorphy reflects time since cave colonization by a lineage. However, this assumption remains to be explicitly tested in North American cave trechines and most other cave taxa. Many previous authors have favored the multiple origins hypothesis whereby once populations became restricted to subterranean habitats, subterranean dispersal was extremely limited except in areas with more expansive karst and interconnected subterranean habitats (Krekeler 1959; Barr 1967; Barr 1985a, Barr and Holsinger 1985). Moreover, many caves in the APP and ILP contain two or more (up to six in the Mammoth Cave System) species of cave trechine beetles belonging to different species groups, supporting multiple cave colonization events (Niemiller et al. 2021; Ober et al. 2022).

As an extension of the multiple origins hypothesis, Barr (1981, 1985, 2004) incorporated a temporal context for cave colonization whereby ancestral surface species occurring in or near karst areas adopted a deep soil existence during glacial periods of the Pleistocene or earlier followed by colonization events into caves during the interglacial periods of the Pleistocene in response to warming and drying of surface habitats, i.e., the climate-relict hypothesis (Jeannel 1943; Holsinger 1988, 2000; Ashmole 1993). Jeannel (1926, 1927, 1928, 1930) and Barr (1967) also hypothesized climate change after the last glacial maximum during the Pleistocene as a major driver of cave colonization and diversification in cave trechine beetles. However, given the large radiation (>150 taxa) of cave trechine species in North America, Ober et al. (2022) did not believe such a recent timeframe for cave colonization and diversification plausible; rather the authors hypothesized multiple colonization events occurring over several interglacial periods occurring as early as the late Pliocene when many caves in the APP and ILP were forming (Poulson and White 1969; Clark 2001; Shofner et al. 2001) to account for cave trechine diversity observed today. Moreover, the presence of two or more cave trechine species occurring in a single cave or cave system (e.g., the Mammoth Cave assemblage contains six cave trechine species; Niemiller et al. 2021) suggests multiple cave colonization events occurred over multiple interglacial periods.

Although taxonomic sampling is incomplete, our analyses support a multiple origin scenario that predates the Pleistocene. Diversification of North American cave trechines began in the middle Miocene with several species groups present by the end of the Miocene and further diversification into the Pliocene, rejecting the Pleistocene climate-relict hypothesis (Jeannel 1943; Barr 1967; Holsinger 1988; 2000; Ashmole 1993) as the primary driver of diversification. Under the climate-relict hypothesis, we might expect simultaneous independent cave colonization events by a single widespread species or multiple closely related species reflected as a polytomy on the inferred species tree and burst of accumulation of lineages on a lineage-through-time plot. In North American cave trechines, we instead see an increase of diversification several million years earlier in the late Miocene into the early Pliocene (Figure 5). Our results are consistent with other recent studies of cave-dwelling beetles. Faille et al. (2011) estimated that a lineage of European trechines colonized caves about 10 Mya, while Ribera et al. (2010) estimated that major lineages of Western Mediterranean cave beetles in the family Leiodidae diverged about 30 Mya. Among other North American cave beetle taxa, diversification events of cave carabid beetles of the genus *Rhadine* (tribe Platynini) in the Edwards Plateau and Balcones Escarpment karst region of Texas occurred within the past 4–5 million years (Gómez et al. 2016) coinciding with a period of cave development in the Balcones Escarpment (Ward 2006; White et al. 2009). Leray et al. (2019) examined diversification of the *hirtus*-group of the small carrion beetle genus *Ptomaphagus* (family Leiodidae), which consists of 19 cave and soil-dwelling species in the central and southeastern United States and co-occurs with trechine cave beetles in many cave systems in the ILP and southern APP. Two main periods of diversification in troglobiotic *Ptomaphagus* were identified: 1) seven geographically distinct lineages diverged across ILP and southern APP 6–8.5 Mya; and 2) a lineage in the southern Cumberland Plateau of Alabama and Tennessee diversified into 12 species over the last 6 million years. Estimated dates of diversification in *Ptomphagus* are quite similar to those in cave trechines. Although significant diversification predates the Pleistocene Epoch, Pleistocene glacial cycles likely have had important impacts on the evolutionary history and biogeography of North American cave trechines. The strong palaeogeographical signal in the distribution of the cave trechine species is likely to be related to their ecological habitats. There are likely to have been multiple independent colonizations, with each lineage having different degrees of morphological adaptation to the subterranean environment. The strongest factor driving the subterranean colonizations may have been the aridification of the climate since the late Miocene (Krijgsman et al. 2000; Micheels et al. 2009). The multiple origins of the subterranean populations or species within the cave trechine group confirmed both by our phylogenetic results is in contrast to hypotheses proposed for other radiations of subterranean beetles, for example in the Pyrenees (Faille et al. 2010, 2013; Ribera et al. 2010; Rizzo et al. 2013; Cieslak et al. 2014), where the entire lineages of species are found exclusively within the deep subterranean environment without morphological variation in some troglomorphic characters (Jeannel 1924, 1928; Salgado et al. 2008). This is likely due to the unique combination of widespread ancestral epigean species with multiple colonizations of the subterranean environment, giving rise to troglobiotic species with very limited geographical ranges that have persisted unaltered over long evolutionary periods.

The hypothesized limited dispersal ability of North American cave trechine beetles (Barr 1967a). Barr 1981; Barr 1985a) and results of this study suggest that troglomorphy evolved in this group multiple times through convergent evolution. In cave trechine evolution, a single cave colonization event would result in a molecular signature of shared loss-of-function (LOF) mutations particularly in loci involved in the regression of eyes and pigmentation among cave trechine lineages (assuming lack of strong selection on particular LOF mutations). In contrast, shared LOF mutations in eye and pigmentation loci are not expected among the cave trechine lineages under a multiple independent cave colonization scenario. However, genomic analyses will be required to test these hypotheses and determine if identical LOF mutations occur in geographically separated cave trechine lineages (single cave invasion subsequently followed by long-distance dispersal) or different LOF mutations occur in geographically distinct lineages (multiple cave invasions subsequently followed by isolation and vicariance). Previous studies supporting multiple independent cave colonization hypothesis include subterranean diving water beetles with distinct mutations in pigmentation and eye opsin genes (Leys et al. 2005; Tierney et al. 2015) and also in the eye rhodopsin locus of geographically distinct lineages of amblyopsid cavefishes in eastern North America (Niemiller et al. 2013).

A single cave colonization scenario requires cave trechines to be able to disperse several kilometers across non-karst terrain that now separates the distributions of many species and species groups. Although dispersal remains to be rigorously examined in cave trechines, it is thought their dispersal ability is quite limited. All cave trechine species are small and wingless, and all but one species has been observed in surface habitats (Ober et al. 2022). Carbonate strata in APP is patchy and discontinuous, and caves are generally smaller and more isolated within this fragmented karst (Hack 1969; Barr 1967a; Barr 1981; Culver 1982; Barr 1985a). Limestone valleys are separated by sandstone ridges in the faulted and folded strata limiting dispersal of cave trechines (Barr 1985). Consequently, most *Pseudanophthalmus* species in the APP are frequently limited to a single or few isolated cave systems (Barr 1965, 1981, 2004; Malabad et al. 2021). For example, 22 of the 64 described *Pseudanophthalmus* species from the APP are single-cave endemics. While evidence suggests a multiple cave colonization scenario is more likely with limited subterranean dispersal, long-distance dispersal cannot be completely ruled out, as long-distance dispersal has been hypothesized for troglobiotic leiodids in central Pyrenees (Rizzo et al. 2013).

In contrast to the APP, only 12 of 84 species in the ILP are endemic to single caves and species generally have larger distributions (Ober et al. 2022). For example, *Darlingtonea kentuckensis* is distributed over an area of ca. 3,728 km^2^ along the western escarpment of the Cumberland Plateau in southeastern Kentucky and adjacent northern Tennessee. Karst in the ILP is characterized by expansive exposures of highly soluble, horizontal-bedded limestones with numerous sinuous branch-like caves systems with four major subregions (Barr 1967): (1) the Bluegrass Region in Kentucky; (2) the western escarpment of the Cumberland Plateau from northeastern Kentucky to northern Alabama; (3) the Central Basin in Tennessee; and (4) the Pennyroyal Plateau from southern Indiana near Bloomington south extending westward into Kentucky and north-central Tennessee. Among these subregions, escarpments of the Pennyroyal Plateau, Central Basin, and the western escarpment of the Cumberland Plateau possess numerous caves that may facilitate dispersal of terrestrial troglobionts, including cave trechines (Barr 1985a). Cave systems in the ILP are more highly connected than in the APP apart from the Bluegrass Region in northern Kentucky near the Ohio River, which is comprised of smaller and more isolated caves that may limit subterranean dispersal in this region (Barr 1967). Cave trechine species are more likely to occur in sympatry in the ILP than in the APP, which may reflect different species colonizing subterranean habitats during different time periods and subsequently dispersing through the highly connected karst of the ILP (Barr 1967).

Although there is evidence for subterranean dispersal in the ILP, vicariance appears to have had a significant role in the diversification and shaping the distributions of many species and species groups in not only the ILP but also the APP. Both hydrological and geological barriers separate both species and species groups in the ILP. For example, within the *P. tenuis* species group *P. barberi* in northern Kentucky is separated from the other five species in the species group by the Ohio River (Barr 2004; Ober et al. 2022). Likewise, two species of the *P. barri* group, *P. barri* and *P. troglodytes*, occur on opposite sides of the Ohio River (Barr 2004). The Ohio River, which formed 0.8 Mya thousand years ago and has down cut via erosion through the cave-bearing strata (Gray 1991; Teller and Goldthwait 1991), appears to be a significant barrier to dispersal for not only troglobionts but also stygobionts (Niemiller et al. 2013). Similarly, the Cumberland River separates the two species of *Nelsonites* with *N. jonesi* to the north and *N. walteri* to the south of the river (Barr 1985a). In contrast, smaller tributaries may not be substantial barriers for all cave trechines, as some species, such as *Neaphaenops tellkampfii* along the Green River in Kentucky and *P. ciliaris* and *P. loganensis* along the Red River in Kentucky and Tennessee, occur on both sides of smaller river and streams (Barr and Peck 1965; Ober et al. 2022). Barr and Peck (1965) hypothesized that flooding may wash beetles out of caves and transporting them downstream to other cave systems on either bank. Barr (1985a) offered support for this hypothesis by examining the relationship between the meander frequencies of rivers and the frequency of cave trechine species occurring on opposite sides of a river. He concluded that the higher the river meander frequency, the more often troglobionts occurred on both sides of the river, suggestive of passive dispersal via flooding.

Barr and Holsinger (1985) hypothesized that dispersal and gene flow between populations cave-dwelling species, particularly troglobionts, can be reduced and ultimately isolated by erosion (i.e., downcutting) of surface water courses into and through cave-bearing strata during the Pliocene and Pleistocene. This vicariance-by-erosion model (sensu Leray et al. 2019) may be a particularly attractive hypothesis for explaining diversification and distribution patterns among taxa within species groups along the Cumberland Plateau and other prominent escarpments of the ILP and also in the APP whereby incisional history and hydrological drainage reorganizations may have influenced the evolutionary history of troglobionts. For example, Leray et al. (2019) propose that the vicariance by erosion model best explained diversification and distributional patterns of *Ptomaphagus* cave fungus beetles, which cooccur with *Pseudanophthalmus*, in the highly dissected southern Cumberland Plateau. The earliest *Ptomaphagus* lineages to divergence in the southern Cumberland Plateau are found in the most peripheral and isolated escarpments, whereas lineages that diverged later are found toward the central region of the southern Cumberland Plateau, a pattern consistent with the predictions of the model.

The biogeography of cave trechine beetles in North America highlights the complex evolution of the eastern North America cave biodiversity hotspots. While our understanding of the evolutionary history and biogeography of this diverse assemblage of cave beetles is incomplete due to incomplete taxonomic sampling, we briefly summarize the biogeography and evolutionary history of eastern Northern American cave trechines based on evidence to date. The surface ancestor(s) to cave trechines appears to have colonized the eastern North America karst regions from an east to west manner during the middle Miocene and Pliocene. The ancestral surface origin of the cave trechine beetles is uncertain but appears to have been in the southern Appalachians. Based on the concatenated phylogeny, the earliest ancestor was likely distributed in the APP and dispersed into the ILP about 11.5 Mya. A surface ancestor likely dispersed into the Ozarks Highlands from the APP between 4.1–7.2 Mya. The APP appears to have served both as a cradle for in situ diversification and as bridge linking the southern Appalachians and ILP, enabling the dispersal and subsequent diversification of these cave beetles. In the APP, there was a burst of diversification in the early Pliocene 7.0 Mya as well as in the Pleistocene 1.2 – 3.7 Mya. After colonization into the ILP, there was further diversification around 10.7 Mya followed by a burst of diversification in the late Miocene 8.0 Mya as well as in the Pliocene into the Pleistocene 1.2 – 5.7 Mya. Based on the dating of radioactive cave sediments, the oldest caves along the western escarpment of the Cumberland Plateau in the ILP are estimated to be 3.5–5.7 million years old (Sasowsky et al. 1995; Anthony and Granger 2004, 2007), however, cave development may have begun much earlier (White 2009).

### Systematics of North American cave trechine beetles

Large (7 mm), slender, morphologically similar species, *Neaphaenops tellkampfii*, *Darlingtonea kentuckensis*, *Ameroduvalius jeanneli*, and *Nelsonites walteri* have been thought to be closely related to *Pseudanophthalmus* species (Valentine 1952; Barr 1972, 1980, 1981, 1985b; Maddison et al. 2019). However, the phylogenetic relationships among these five genera have been unclear due to limited sampling of species within the genus *Pseudanophthalmus*. We found that *Pseudanophthalmus* as currently recognized is paraphyletic with respect to the four other genera (*Neaphanops*, *Darlingtonea*, *Nelsonites*, and *Ameroduvalius*) suggesting that these genera are derived from *Pseudanophthalmus*. The relatively widespread distribution of *Neaphaenops tellkampfii* includes the Pennyroyal Plateau in Kentucky and the distributions of the other three genera includes the western escarpment of the Cumberland Plateau in the ILP of eastern Kentucky and north-central Tennessee. The distributions of all four genera overlap with *Pseudanophthalmus*, and they often co-occur within the same cave systems (Ober et al. 2022).

Contrastingly, *Xenotrechus*, which is known from a few caves in southeastern Missouri west of the Mississippi River (Barr and Krekeler 1967) is believed to be distantly related to *Pseudanophthalmus* and trechine genera. We were unable to include any *Xenotrechus* species in our study. Although the phylogenetic placement of genus *Xenotrechus* is unknown, *Xenotrechus* is hypothesized to be related to the eastern European genera *Chaetoduvalius* and *Geotrechus* lacking any close relatives in North America (Barr and Krekeler 1967). At present, we assume the exclusion of *Xenotrechus* has minimal impacts on our biogeographic interpretations.

While the phylogeny of cave trechines in eastern North America is incomplete, Barr’s (2004) species group classification arrangement provides a framework for how several species of cave trechines may be related. Barr categorized species into species groups based on shared morphological characters, such as the shape of a groove at the apex of the elytra and features of the male genitalia, and also their distributions in karst regions. Barr (1981, 1985a) hypothesized that closely related species morphologically generally co-occur in the same geographical area.

Our results showed that apart from the *cumberlandus*, *grandis*, *jonesi*, and *simplex* species groups, all other species groups proposed by Barr (2004) formed monophyletic groups. For the species groups that were not recovered as monophyletic, morphological similarity may reflect morphological convergence and cryptic speciation, which is frequently reported in phylogenetic studies of cave organisms (Wiens et al. 2003; Derkarabetian et al. 2010; Niemiller et al. 2012; Maddison et al. 2019). However, additional and more comprehensive taxonomic sampling is warranted.

## Conclusions

This study represents the first effort to establish a time-calibrated phylogenomic framework for cave trechine beetles in North America, elucidating a rich and intriguing history of evolution. Our results conflicted with previous generic and many species-group taxonomic classification hypotheses based on morphology. In particular, the genera *Neaphanops*, *Darlingtonea*, *Nelsonites*, and *Ameroduvalius* were nested within *Pseudanopthalmus* and some species groups were not recovered as monophyletic. The surface ancestor of cave trechines likely began colonizing caves in the Appalachians Ridge and Valley in the middle Miocene around 11.5 Mya. The evolution of *Pseudonaphthalmus* is characterized by rapid early radiation followed by a series of dispersal events into the Ozark Highlands and Interior Low Plateau, with many of the major clades attaining their present-day geographic distributions by the early Miocene and with multiple additional episodes of cave colonization and diversification occurring throughout the Pliocene and Pleistocene. Additional research is needed to better characterize the levels of diversity, speciation, and the origins of cave trechines and to understand their phylogeographic patterns in eastern North America. In summary, molecular systematics and biogeography of these unique cave beetles offer a model for other comparative evolutionary and ecological studies of troglobionts to further our understanding of factors driving speciation and biogeographic patterns.

## Figures and tables

## Supplemental figures and tables

**Supplemental Figure 1.**
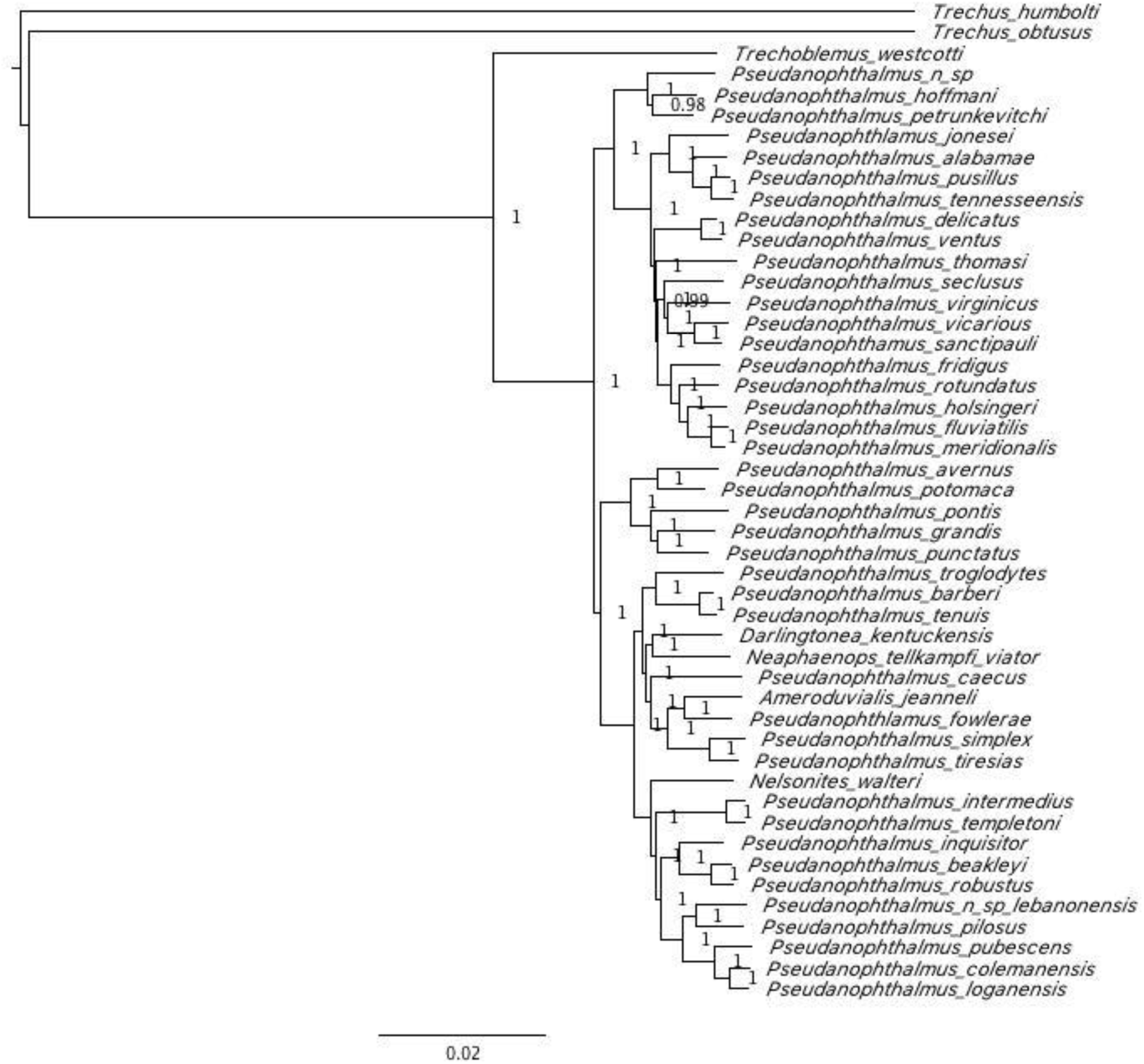
Bayesian phylogeny of cave trechine beetles from eastern North America inferred from 50% complete concatenated UCE matrix. Numbers indicate support values (Bayesian posterior probability) for all nodes.

**Supplemental Figure 2.**
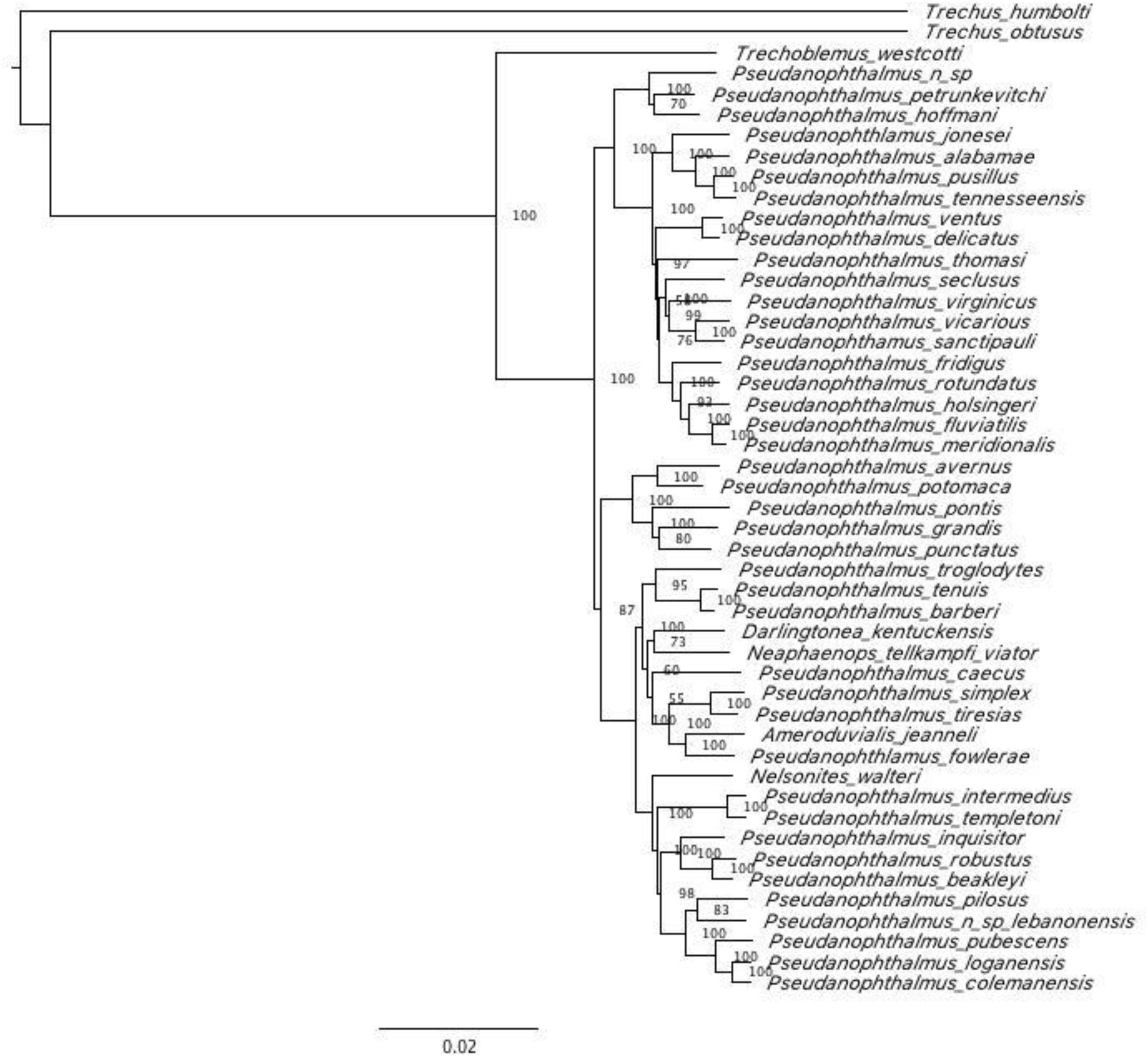
Maximum-likelihood phylogeny of cave trechine beetles from eastern North America inferred from 50% complete concatenated UCE matrix. Numbers indicate support values (maximum-likelihood) for all nodes.

**Supplemental Figure 3.**
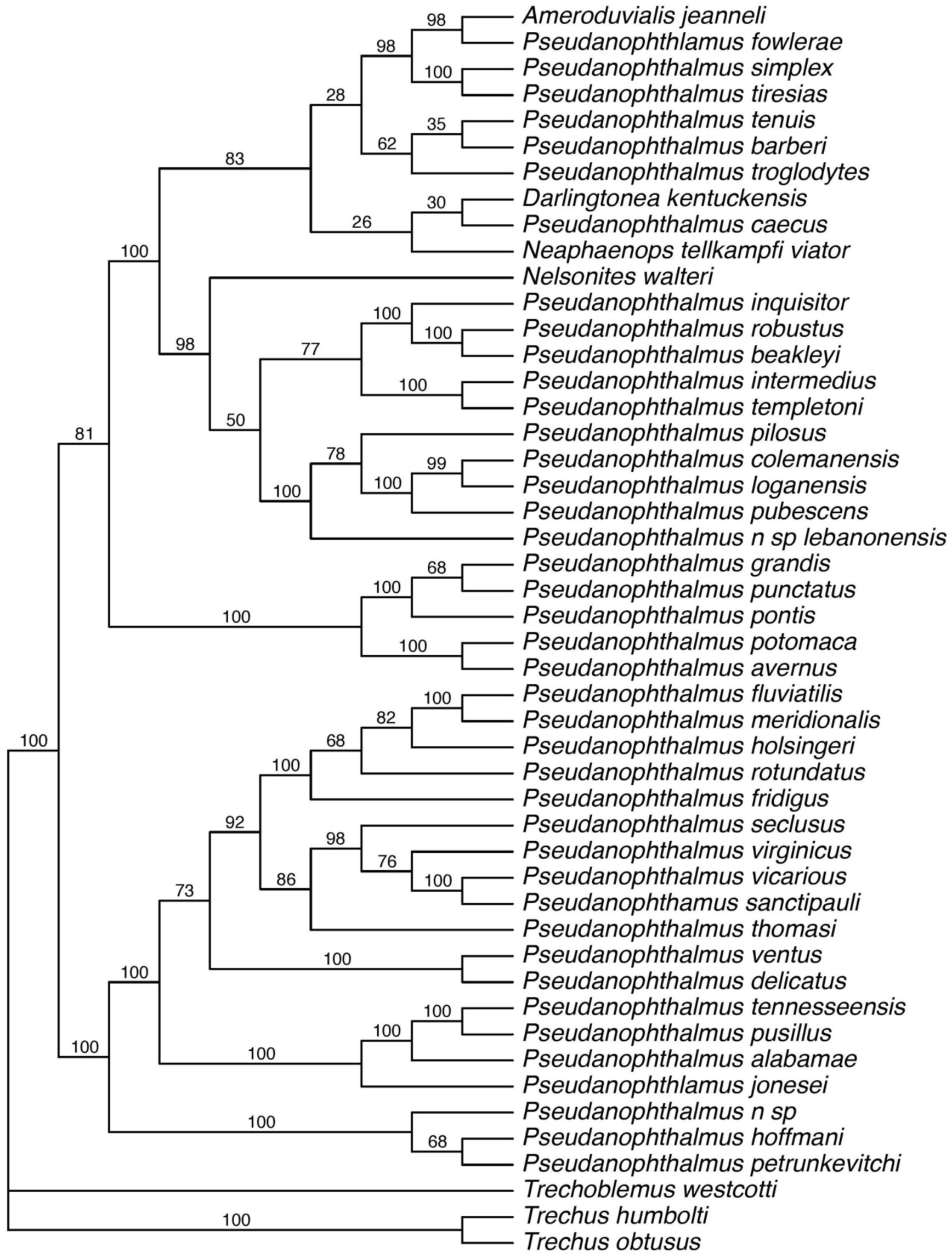
Phylogenetic relationships among the cave trechine beetles based on SVDQuartets coalescent species trees with 50% majority rule consensus for SVDQuartets. Node values indicate bootstrap support values.

**Supplemental Figure 4.**
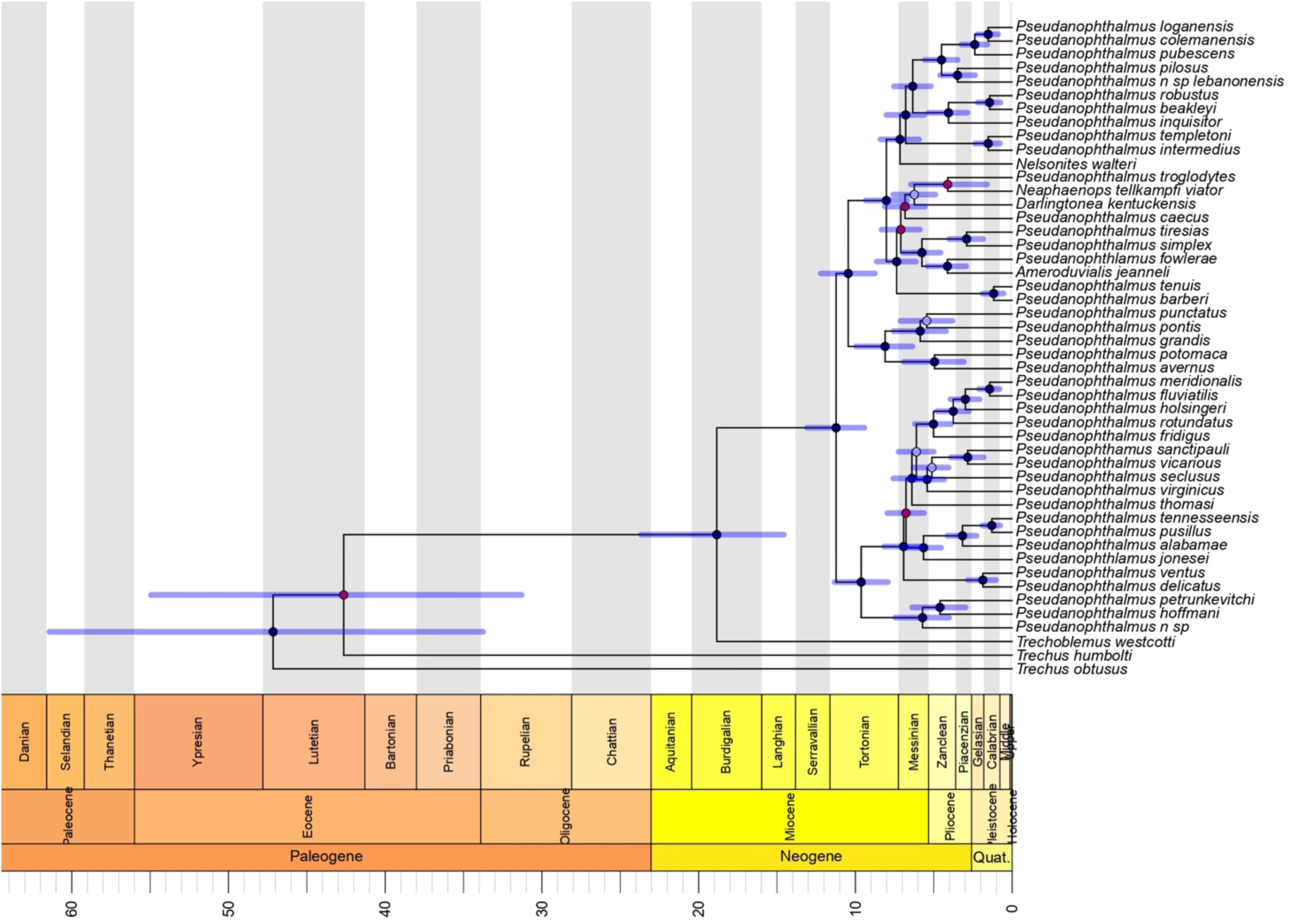
Time-calibrated maximum clade credibility tree with *Trechus* outgroup inferred from 75% complete concatenated UCE matrix, summarized by TreeAnnotator, and plotted with a geological time scale using the strap package in R. Phylogeny dated using a Bayesian relaxed clock method in BEAST. Branches are proportional to time in millions of years. The 95% confidence intervals for the ages of basal branches in the tree are indicated with transparent blue bars. The internal nodes of the tree are indicated with circles, where circles mark nodes with posterior probability:

**Supplemental Figure 5.**
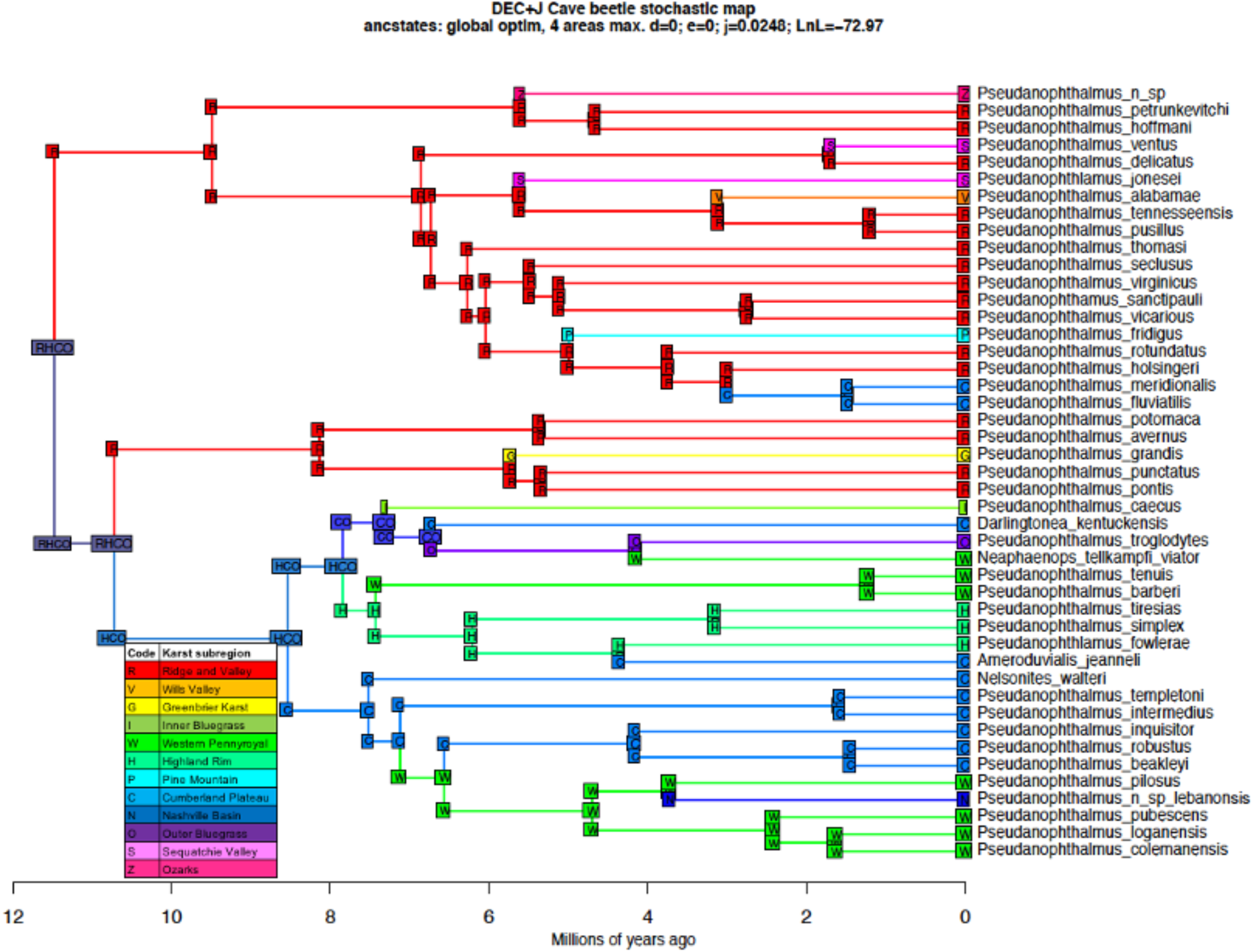
Ancestral area estimation for cave trechine beetles from eastern North America based on the preferred DEC+J model. Ancestral areas were estimated across the time-calibrated phylogeny inferred from 75% complete concatenated UCE matrix. Most probable ancestral karst sub region range at each node shown. Corner positions represent the geographic karst sub region range immediately after a cladogenetic event.

**Supplementary Table 1.** Information on specimen vouchers, sample, data yield (Mb), raw Illumina reads before and after quality filtering and trimming, and SRA accession numbers

| Voucher | Species | Yield (Mb) | Raw Reads | Raw Reads<br>after QC | SRA accession |
| --- | --- | --- | --- | --- | --- |
| DNA240 | *Nelsonites walteri* | 389 | 1290679 | 1264267 | SAMN31468728 |
| DNA257 | *Pseudanophthalmus pubescens* | 751 | 2489047 | 2433261 | SAMN31468729 |
| DNA289 | *Pseudanophthalmus n. sp. lebanonensis* | 1014 | 3359680 | 3284768 | SAMN31468730 |
| DNA296 | *Ameroduvialis jeanneli* | 1433 | 4746236 | 4661544 | SAMN31468731 |
| DNA303 | *Pseudanophthalmus meridionalis* | 1185 | 3926710 | 3849578 | SAMN31468732 |
| DNA313 | *Pseudanophthalmus tennesseensis* | 2648 | 8770180 | 8649185 | SAMN31468733 |
| DNA315 | *Pseudanophthalmus robustus* | 1218 | 4034493 | 3961570 | SAMN31468734 |
| DNA370 | *Pseudanophthalmus colemanensis* | 2234 | 7400154 | 7291619 | SAMN31468735 |
| DNA385 | *Pseudanophthalmus fluviatilis* | 963 | 3190448 | 3123659 | SAMN31468736 |
| DNA404 | *Pseudanophthalmus jonesei* | 1574 | 5213092 | 5133024 | SAMN31468737 |
| DNA410 | *Pseudanophthalmus fowlerae* | 2701 | 8944267 | 8824422 | SAMN31468738 |
| DNA467 | *Darlingtonia kentuckensis* | 1903 | 6304505 | 6211516 | SAMN31468739 |
| DNA474 | *Pseudanophthalmus intermedius* | 1545 | 5118836 | 5036087 | SAMN31468740 |
| DNA492 | *Pseudanophthalmus delicatus* | 1263 | 4185208 | 4120192 | SAMN31468741 |
| DNA498 | *Pseudanophthalmus loganensis* | 1988 | 6585439 | 6500856 | SAMN31468742 |
| DNA507 | *Pseudanophthalmus avernus* | 2269 | 7515566 | 7403566 | SAMN31468743 |
| DNA510 | *Pseudanophthalmus tiresias* | 890 | 2949246 | 2902518 | SAMN31468744 |
| DNA511 | *Pseudanophthalmus beakleyi* | 2132 | 7061910 | 6957591 | SAMN31468745 |
| DNA525 | *Pseudanophthalmus tenuis* | 1982 | 6564135 | 6469009 | SAMN31468746 |
| DNA530 | *Pseudanophthalmus pusillus* | 3450 | 11426668 | 11283637 | SAMN31468747 |
| DNA538 | *Pseudanophthalmus grandis* | 991 | 3283450 | 3236038 | SAMN31468748 |
| DNA541 | *Pseudanophthalmus fridigus* | 330 | 1093094 | 1071502 | SAMN31468749 |
| DNA544 | *Pseudanophthalmus troglodytes* | 46 | 154671 | 146144 | SAMN31468750 |
| DNA554 | *Pseudanophthalmus caecus* | 136 | 453183 | 441279 | SAMN31468751 |
| DNA570 | *Pseudanophthalmus ventus* | 1230 | 4074848 | 4022862 | SAMN31468752 |
| DNA571 | *Pseudanophthalmus templetoni* | 1811 | 5997310 | 5918933 | SAMN31468753 |
| DNA574 | *Pseudanophthalmus inquisitor* | 850 | 2814717 | 2764858 | SAMN31468754 |
| DNA577 | *Pseudanophthalmus pontis* | 1493 | 4944655 | 4875265 | SAMN31468755 |
| DNA582 | *Pseudanophthalmus punctatus* | 2048 | 6783111 | 6676826 | SAMN31468756 |
| DNA593 | *Pseudanophthalmus rotundatus* | 2839 | 9401238 | 9258512 | SAMN31468757 |
| DNA600 | *Pseudanophthalmus simplex* | 1637 | 5423131 | 5334661 | SAMN31468758 |
| DNA607 | *Pseudanophthalmus barberi* | 1530 | 5068330 | 4998957 | SAMN31468759 |
| DNA618 | *Pseudanophthalmus pilosus* | 1556 | 5153059 | 5085574 | SAMN31468760 |
| DNA631 | *Pseudanophthalmus holsingeri* | 2181 | 7222348 | 7103880 | SAMN31468761 |
| DNA632 | *Pseudanophthalmus thomasi* | 1655 | 5482482 | 5396298 | SAMN31468762 |
| DNA656 | *Neaphaenops tellkampfi viator* | 1400 | 4636586 | 4532101 | SAMN31468763 |
| DNA670 | *Pseudanophthalmus seclusus* | 1522 | 5040936 | 4962632 | SAMN31468764 |
| DNA685 | *Pseudanophthalmus virginicus* | 2571 | 8516008 | 8386082 | SAMN31468765 |
| DNA689 | *Pseudanophthalmus n.sp.* | 1387 | 4593760 | 4526457 | SAMN31468766 |
| DNA748 | *Pseudanophthalmus alabamae* | 1927 | 6383048 | 6290294 | SAMN31468767 |
| DNA756 | *Pseudanophthalmus potomaca* | 1799 | 5959574 | 5849553 | SAMN31468768 |
| DNA758 | *Pseudanophthamus sanctipauli* | 2085 | 6904911 | 6799444 | SAMN31468769 |
| DNA766 | *Pseudanophthalmus vicarious* | 1992 | 6598288 | 6498008 | SAMN31468770 |
| DNA767 | *Pseudanophthalmus petrunkevitchi* | 1955 | 6473609 | 6386504 | SAMN31468771 |
| DNA781 | *Pseudanophthalmus hoffmani* | 1858 | 6154080 | 6053781 | SAMN31468772 |
| DNA786 | *Trechoblemus westcotti* | 1103 | 3652403 | 3583207 | SAMN31468773 |
| DNA787 | *Trechus obtusus* | 108 | 358362 | 342995 | SAMN31468774 |
| DNA788 | *Trechus humboldti* | 591 | 1957105 | 1900383 | SAMN31468775 |

**Supplementary Table 2.** Comparison of dispersal-extinction-cladogenesis (DEC) models with jump dispersal (+J) and without (+J) for cave trechine beetles based on their dispersal within karst sub region. Abbreviations as follows: LnL, loglikelihood; numparams, number of parameters in each model; d, dispersal rate; e, extinction rate; j, founder-event speciation rate; AIC, Akaike Information Criterion; LRT, likelihood-ratio test.

| Model | LnL | numparams | d | e | j | AIC | AIC_wt | LRT<br>pval |
| --- | --- | --- | --- | --- | --- | --- | --- | --- |
| DEC | -101.8 | 2 | 0.0048 | 0.026 | 0 | 207.6 | 8.00E-13 |  |
| DEC+J | -72.97 | 3 | 1.00E-12 | 1.00E-12 | 0.025 | 151.9 | 1 | 3.10E-14 |

